# Preserved cerebellar functions despite structural degeneration in older adults

**DOI:** 10.1101/2025.06.18.660418

**Authors:** Anda de Witte, Anouck Matthijs, Benjamin Parrell, Dante Mantini, Jolien Gooijers, Jean-Jacques Orban de Xivry

## Abstract

Aging is frequently perceived negatively due to its association with the decline of various brain and bodily functions. While it is evident that motor abilities deteriorate with age, it is incorrect to assume that all aspects of movement execution are equally affected. The cerebellum, a brain region that is closely involved in motor control among other functions, undergoes clear structural changes with aging. While several studies suggest that cerebellar degeneration causes age-related motor control deficits, other studies suggest that the cerebellum might act as a motor reserve and compensate for its structural degeneration, leaving cerebellar motor function intact despite cerebellar degeneration. The present study aims at thoroughly investigating the impact of age on cerebellar function across an array of tasks and domains.

We investigated cerebellar motor and cognitive functions across the lifespan by examining 50 young adults (20–35 years), 80 older adults (55–70 years), and 30 older-old adults (>80 years). Participants completed a test battery comprising seven motor control tasks and one cognitive task, each designed to probe cerebellar function through different paradigms. This multi-task approach allowed for a comprehensive evaluation of performance patterns, providing a balanced perspective on cerebellar function across the different age groups. In addition, we analyzed outcomes from the same tasks that, while related to movement, were not specifically linked to cerebellar function. Structural magnetic resonance imaging was also conducted to assess whether cerebellar atrophy was present in the older and older-old groups compared to the young.

Our results revealed that, despite age-related cerebellar degeneration, cerebellar functions in older adults remained intact compared to young adults, even in adults above 80 years old. In contrast, the sensorimotor measures that were not directly linked to cerebellar function exhibited a clear pattern of decline in older adults, and were further deteriorated in the older-old adults compared to the older adults.

These findings indicate that cerebellar motor control functions remain largely preserved with age, providing compelling evidence that the cerebellum possesses a remarkable degree of functional resilience and redundancy. This suggests that cerebellar circuits may be uniquely equipped to preserve function despite structural degeneration.

## Introduction

Healthy older adults often experience being negatively judged due to age-based stereotypes (Barber, 2020), as aging is frequently associated with a growing number of functional impairments (Chen et al., 2023). Mobility impairments are particularly common and raise significant concern, as they compromise individuals’ independence and often lead to increased reliance on care from others (Ferrucci et al., 2016). These impairments are driven by multiple factors, including declines in muscle strength, sensory deterioration such as hearing loss and visual impairments, neurodegenerative changes, and other age-related conditions that affect balance, coordination, and motor control (Cavallari et al., 2013; Ferrucci et al., 2016). While extensive research has been dedicated to developing therapies and medications to mitigate or prevent these deficits (Izquierdo et al., 2021; C.-C. Wu et al., 2022), it is equally important to gain a deeper understanding of the specific contributions of individual mechanisms to motor behavior in older people. To do so, we must not only examine the mechanisms that decline with age, but also recognize and highlight those that remain stable, or may even improve at older age (Poirier et al., 2021). This balanced perspective can inform more effective interventions to support healthy aging, while also promoting a more positive perception of aging, further reinforcing the pursuit of well-being in later life (Barber, 2020; Feliciano et al., 2022).

In this context, we aimed to investigate how the cerebellum functions in older age. The cerebellum has traditionally been recognized for its role in movement control (Paulin, 1993), but accumulating research indicates that it is involvement in a wider range of functions as well, including cognition, timing, and emotion (Habas et al., 2009; Manto et al., 2012; Prati et al., 2024; Strick et al., 2009). Studies of patients with cerebellar damage have significantly advanced our understanding of the cerebellum’s functions, demonstrating that these individuals often experience inaccurate and clumsy limb movements, postural and gait instability, nystagmus, and speech difficulties. These impairments primarily arise from disruptions in cerebellar timing (Ivry & Keele, 1989) and coordination mechanism (Matsuo et al., 2005), particularly those involved in acquiring sensory information and synchronizing it with motor predictions (Manto et al., 2012; Statton et al., 2018). Timing and coordination deficits are evident in patients with impaired intra-limb coordination, where the execution of smooth and precise movements is disrupted (Matsuo et al., 2005). Moreover, impaired timing abilities lead to inaccurate motor responses due to diminished capacity to anticipate self-generated actions (Bruttini et al., 2015), such as insufficient grip force anticipation during object lifting (Rost et al., 2005), and delayed anticipatory eye movements (Lekwuwa et al., 1995). Patients with cerebellar damage also showed reduced implicit adaptation, struggling to adjust their movements in response to external perturbations in multiple motor domains such as reaching movements, eye movements and speech (Golla et al., 2008; Morehead et al., 2017; Parrell et al., 2017). Beyond the motor domain, cerebellar damage affects cognitive and affective processes as well, such as impairments in mental rotation tasks (McDougle et al., 2022) and disruptions in emotional learning such as fear conditioning (Hwang et al., 2022).

When it comes to aging, the cerebellum is one of the brain structures identified as particularly vulnerable to age-related deterioration. Cerebellar gray matter loss begins as early as around the age of 50 and accelerates exponentially thereafter, resulting in more severe cerebellar gray matter loss in older age (Luft, 1999; Terribilli et al., 2011; Walhovd et al., 2005). The loss of cerebellar white matter volume occurs at a later age but progresses more extensively than cerebellar gray matter loss (Hoogendam et al., 2012). Since age-related cerebellar degeneration follows a pattern similar to, though less severe than, that seen in patients with cerebellar disease (Hulst et al., 2015), it raises the question of how it impacts the motor function in older adults.

Currently, inconsistencies exist about whether behavioral cerebellar deficits occur and if they affect movement control in the elderly. A prevailing theory is that age-related degeneration of the cerebellum contributes directly to declines in various functions in older adults, including executive functioning, working memory, balance, and proprioception (Bernard & Seidler, 2013; Boisgontier & Nougier, 2013; Droby et al., 2021; Liang & Carlson, 2020; Raz et al., 2000; Rosano et al., 2007). However, much of the evidence supporting this theory comes from studies that narrowly focused on a single task (Boisgontier & Nougier, 2013; Miller et al., 2013; Woodruff-Pak et al., 2001) or on assessments within similar cerebellar domains such as balance and gait (Droby et al., 2021; Rosano et al., 2007), instead of capturing a broader range of specific cerebellar functions. Isolating the specific contribution of the cerebellum to task performance is tricky, as any task will typically engage a broad network involving cortical ad subcortical areas beyond the cerebellum (Caligiore et al., 2017; Prati et al., 2024). Therefore, accurately assessing cerebellar function requires carefully designed tasks that specifically isolate cerebellar contributions. As assessing cerebellar functioning inherently depends on both the quality of cortical inputs and the effectiveness of cerebellar feedback to the cortex, it is difficult to isolate cerebellar function from these other factors (Bernard, 2022; Bernard & Seidler, 2014). Relying solely on daily-life symptoms to assess cerebellar patients is often too unspecific, as many of the symptoms overlap with those of other patient groups, suggesting the influence of extra-cerebellar factors. For example gait and balance problems, as well as motor timing problems occur in Parkinson’s patients as well (Horak & Mancini, 2013; Jones & Jahanshahi, 2014). Therefore, we emphasize the importance of assessing cerebellar functions using insights from previous studies on cerebellar patients that carefully distinguished and identified the factors affecting task outcomes and accordingly select tasks that allow for such distinctions. Based on the literature on these tasks, many cerebellar functions that are impaired in cerebellar patients appear to be preserved in older adults. Examples of preserved cerebellar contributions include intact timing functions, such as rhythmic finger-tapping performance (Duchek et al., 1994) and the preserved ability to act upon temporal predictions about a moving target (Filip et al., 2019). Other studies found that cerebellar predictive functions that enable anticipation of the sensory consequences of one’s own actions remain unaltered by aging as well in several different contexts. For instance, when generating up-and-down movements of an object, older adults produced well-coordinated adjustments in grip force to predictable load force changes (Gilles & Wing, 2003), suggesting intact cerebellum-dependent prediction mechanisms responsible for this coordination (Rost et al., 2005; Ulloa et al., 2003). Similarly, during self-generated touch on their own hand or arm, older adults exhibited greater sensory attenuation compared to younger adults. This suggests an increased reliance on cerebellar motor predictions, possibly compensating for reduced sensitivity in sensory feedback about the generated touch (Parthasharathy et al., 2022; Wolpe et al., 2016). Although, to our knowledge, no sensory attenuation deficits have been reported in cerebellar patients, several studies have pinpointed the cerebellum’s crucial role in attenuating activity within sensory brain regions (Cao et al., 2017; Kilteni et al., 2020; Wolpe et al., 2016). In the domain of motor coordination, older individuals exhibit intact ability to represent and use inter-segmental dynamics for the control of movement (Lee et al., 2007), an ability that is impaired in individuals with cerebellar lesions (Bastian et al., 1996). Further, implicit adaptation in visuomotor arm-reaching tasks remains intact in older adults and may even be enhanced, indicating preserved cerebellar internal model recalibration (Hermans et al., 2025; Van De Plas & Orban de Xivry, 2026; Vandevoorde & Orban De Xivry, 2019).

Taken together, these previous findings suggest that while the cerebellum undergoes structural degeneration with age, this does not necessarily result in a corresponding decline in function. This supports the idea of an age-related brain reserve, reflecting the brain’s resilience and capacity to maintain function despite significant structural changes (Mitoma et al., 2020). Nowadays, the concept of brain reserve is widely used and has been defined in various ways in the literature (Cabeza et al., 2018; Sumowski et al., 2013). It was first introduced in the field of cognition, where some individuals with Alzheimer’s disease pathology exhibited no symptoms or had milder symptoms than expected based on the extent of their brain pathology (Stern, 2002). Similar functional preservation has been observed in patients with Parkinson’s disease, showing that most of them do not do not exhibit symptoms until they have lost 50–70% of the cells in the thalamus and substantia nigra (Fearnley & Lees, 1991). These findings directly challenged the widely held assumption that structural decline inevitably leads to functional deterioration (Hogan, 2004; Persson et al., 2006). One plausible explanation for the existence of such a reserve is the engagement of adaptive mechanisms that compensate for structural loss, thereby preserving function (Habeck et al., 2016). Neuroimaging studies support this by showing that older adults exhibit increased brain activity and enhanced functional connectivity during task performance (Cabeza et al., 2018; Goh & Park, 2009; Mattay et al., 2006). This suggests that increased neuronal activity and the recruitment of additional brain regions serve as compensatory mechanisms for maintaining function (Cabeza et al., 2018; Morcom & Johnson, 2015).

The preserved cerebellar functions in the elderly, combined with emerging reserve theories, led to the idea of a similar reserve for cerebellar functions in the context of age-related cerebellar structural loss (Arleo et al., 2023; Bordignon et al., 2021; Gelfo & Petrosini, 2022; Mitoma et al., 2020). However, while previous literature consistently highlights the strong age-related structural degeneration of the cerebellum and its consequences for cerebellar motor functions (Bernard & Seidler, 2014; Y. Wang et al., 2024), there is actually no clear consensus on whether cerebellar functions are preserved or decline in older adults. This requires a comprehensive test battery that evaluates cerebellar function across a diverse range of tasks, within the same sample. Additionally, this approach requires a large sample size to accurately detect potential differences, along with ensuring that subjects are sufficiently old to exhibit age-related cerebellar function impairments.

In the current study, we aimed to investigate cerebellar motor control function in young, older and older-old (above 80 years old) adults using an extensive test battery. The tasks assessed motor timing, motor coordination, implicit adaptation, and cognition. Task selection and cerebellar outcome measures were primarily based on previous findings of functional deficits in cerebellar patients (Table 1). We hypothesize that older adults will exhibit a generally preserved pattern of cerebellar function across tasks, despite structural cerebellar loss, supporting the notion of a cerebellar reserve. While we caution against overgeneralizing cerebellar function across contexts, we argue that evidence of preserved cerebellar motor function is only convincing when demonstrated across multiple tasks that engage different aspects of cerebellar processing. This is because the cerebellum always operates within networks involving both the body and neocortical areas, making single-task results insufficient to substantiate this claim. Furthermore, we distinguished cerebellar function outcomes from more general sensorimotor outcomes, extracted from the same tasks, to assess the cerebellar function independently of overall task performance. Finally, we used neuroimaging to assess cerebellar gray and white matter volumes to confirm that our older sample exhibited reduced volumes relative to young adults.

**Table 1.**
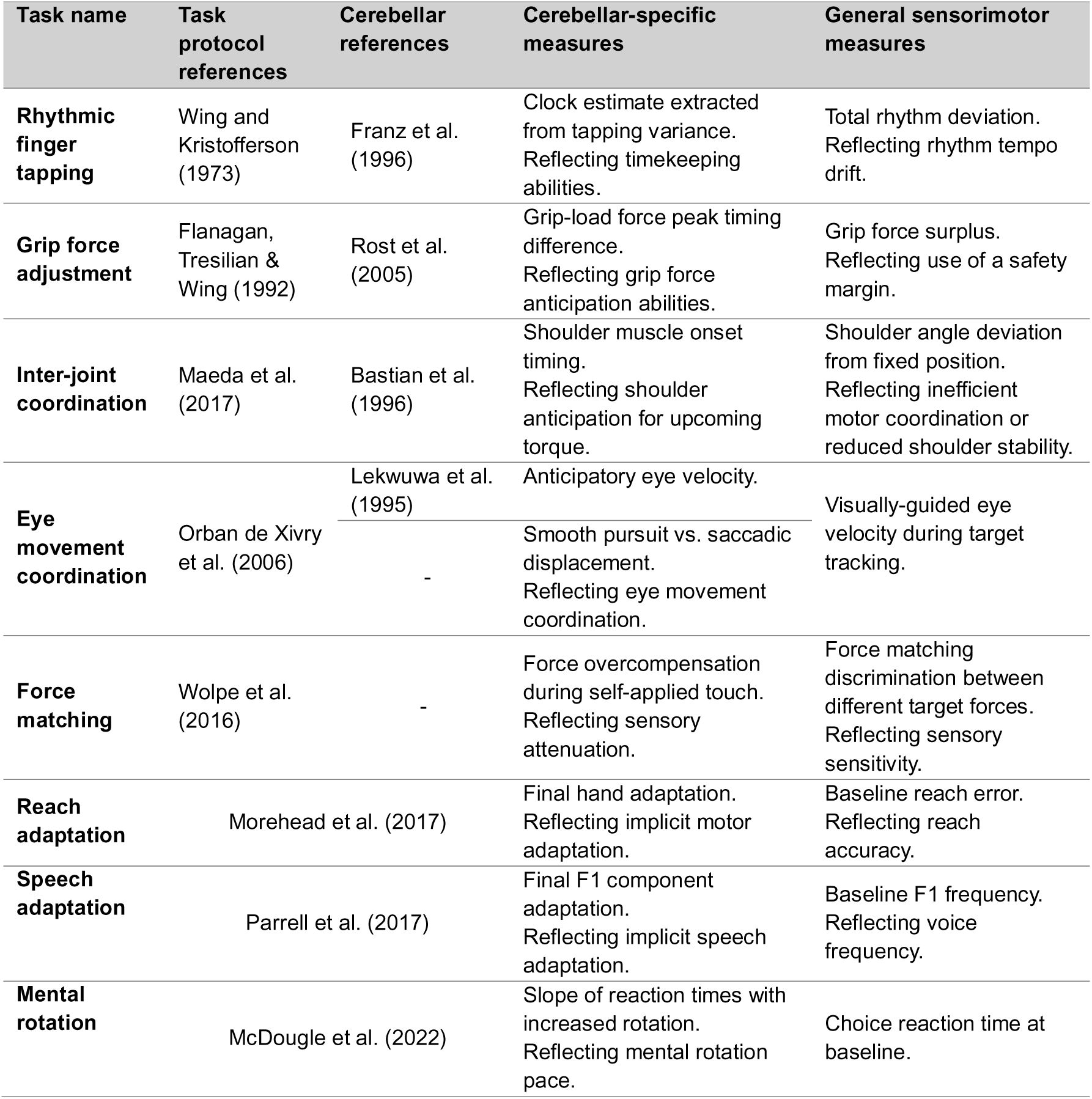
Overview of behavioral test battery.

## Methods

### Participants

In total we recruited 167 healthy adults: 50 young adults between 20 and 25 years (mean age: 23.3 ±2.5, 28 females), 82 older adults between 55 and 70 years (mean age: 63.1 ±4.6, 45 females) and 35 older-old adults above 80 years (mean age: 81.9 ±1.7, 14 females). During recruitment, participants were verbally screened for right-handedness, non-smoker status and good mental and physical health, including being free of neurological diseases. After providing written informed consent, all participants were screened with the Edinburgh handedness inventory (Oldfield 1971) to confirm their right-handedness. We checked cognitive functions of older and older-old participants using the Montreal Cognitive Assessment test (MoCA). We assessed cognitive functioning in both older and older-old participants using the Montreal Cognitive Assessment (MoCA). A minimum score of 23 out of 30 was required for inclusion, following the recommendation by Carson et al. (2018), who demonstrated that this reduced cutoff yields fewer false positives and provides better overall diagnostic accuracy than the original 26/30 threshold. We adopted this criterion to ensure that our sample was not limited to only the highest-performing older adults.

This screening aimed to exclude participants with symptoms of potential mild cognitive impairment, ensuring a sample of cognitively healthy individuals more likely to understand task instructions accurately. None of the task components assessed in the MoCA overlapped with those included in our behavioral test procedure. Six participants were excluded based on MoCA criteria. One participant was left out of the analysis because of repeatedly falling asleep during measurements. The final dataset included a total amount of 50 young adults, 80 older adults and 30 older-old adults. The study was approved by the Ethics Committee Research UZ / KU Leuven (project ID: S66650).

### Testing procedure

This study is part of a larger project that involved five hours of behavioral assessments across two sessions and one hour of MRI scanning. We selected a range of behavioral tasks to assess cerebellar motor and cognitive functions (see description below and Table 1 for an overview). For the older-old participants, tasks were distributed differently between sessions compared to the protocol used for young and older participants, because this group did not perform all tasks or all task levels, mostly for tasks outside the scope of the present study. This adjustment was necessary to ensure equal session lengths. For the task order, we prioritized placing cognitively demanding tasks in the first half of each session and aimed to alternate between physically intensive tasks, attention-demanding tasks, and shorter versus longer tasks. There was no fixed interval between the two behavioral sessions. Ideally, both were scheduled within one week, but in practice, the timing was adapted to participants’ availability. Across all participants, this resulted in a mean inter-session interval of 7.40 days (± 9.03; range = 0-63 days). The average interval between the behavioral sessions and the MRI scanning was 6.86 days (± 8.90; range = 0-83 days).

Task performance outcomes linked to these cerebellar functions were labeled as cerebellar-specific measures. Although these tasks were selected based on their dependence on the cerebellum, we also assessed other measures that strongly contributed to task performance but were not directly related to cerebellar function and labeled them as general sensorimotor measures. The behavioral tests were performed by all three age groups, except for the eye movement task, which was not performed by participants in the older-old group due to the high likelihood of tracking inaccuracies caused by deep-set eyes or drooping eyelids. MRI scanning was also optional for this group, as many participants had metal implants or experienced discomfort, limiting feasibility. Ultimately, we successfully obtained MR images from 20 participants in the older-old group (67%).

### Rhythmic finger tapping

We assessed motor timing abilities based on the rhythmic finger tapping task protocol of Wing and Kristofferson (1973). Participants were seated at a table and used the spacebar of a computer keyboard for tapping. We instructed them to tap along with the rhythm of a given sequence of tones and to continue tapping at the same rhythm after the sequence ended until instructed to stop.

Each trial began with a sequence of tones set at a frequency of 750 Hz, with each tone lasting 150 ms. The inter-stimulus interval was set at 600 ms (corresponding to 1.67 Hz). After 13 tones, the auditory sequence stopped, and participants continued tapping in rhythm for an additional 31 taps (unguided taps). To ensure task comprehension, each participant first completed one practice trial before proceeding to the main task that, which consisted of three test trials. In addition to this task, participants followed a similar procedure for two additional rhythm conditions with lower frequencies. These additional conditions, which explore difficulties in adapting to different rhythms, are examined in a separate project.

We calculated inter-tapping intervals using the final 30 unguided taps from each of the three test trials, with the first unguided tap excluded to avoid potential effects of the guided-to-unguided transition. Trials that contained intervals shorter than 150 ms, classified as accidental double-taps, were excluded from the analysis. Based on this criterion, 14 trials (3%) were removed. We also screened for unusually long intervals, specifically, those exceeding twice the reference interval (>1,200 ms), as potential indicators of participant inattention. However, all such cases occurred in trials that had already been excluded due to double-taps. Participants with fewer than two valid trials remaining were excluded from further analysis; which applied to four individuals. For each of the remaining trials, we computed the average inter-tapping interval. Clock and motor variance estimates were calculated based on the model (Wing & Kristofferson, 1973). Clock variance refers to variability in the internal timekeeping mechanism that regulates the timing of internal triggers to tap. The timekeeping function is theorized to depend on a dedicated timing loop, with the cerebellum as its primary component, a view supported by findings of increased clock variance particularly in patients with cerebellar degeneration or damage (Franz et al., 1996; Harrington, 2003; Ivry & Keele, 1989). For this reason, we selected clock variance as a cerebellar-specific measure. In the same model, motor variance reflects variability in the delay between the internal trigger and the physical execution of each tap, capturing inconsistencies in motor executions. We calculated motor variance to complete all steps of the model; however, it was not included in further analysis, as it was mildly affected in patients with cerebellar disease but more significantly impacted in those with cortical or peripheral damage, making it difficult to pinpoint the underlying mechanism (Ivry & Keele, 1989). For further analysis of the timekeeping mechanism, we selected only the clock delay estimates.

Clock variances (S_C_) were calculated based on the tapping intervals using the following equations:

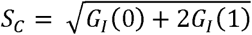

With *G_I_*(1) as the covariance between adjacent inter-tap intervals:

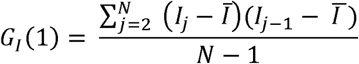

and *G_I_*(0) as total variance of the observable inter-tap intervals:

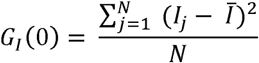

where *I_j_* refers to the j^th^ response interval, and:

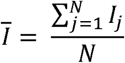

with N = 30.

The Wing and Kristofferson model is based on the idea that variation in the motor mechanism produces a negative correlation between adjacent intervals. For each of our trials, we tested this assumption by calculating whether the lag one serial correlation (P_I_(1)) was negative, using:

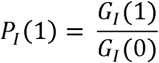

A negative *P_I_*(1) indicates that long intervals tend to be followed by short ones, and short ones, and short intervals by long ones, reflecting negative dependence. The model’s assumptions are violated if *P_I_*(1) = 0, implying statistically independence, or if *P_I_*(1) > 0, indicating that long intervals tend to be followed by long ones and short by short ones.

The assumption of a negative serial correlation was violated in 131 trials (28.17%). In such cases, in accordance with the methods of Ivry and Keele (1989), we set the motor delay estimates to zero, as it is theorized that all the tapping variability then comes from the timekeeper mechanism. This adjustment ensured that clock variance did not exceed total variability. An additional 20 trials were found in which the lag one serial correlation was lower than −0.5 (4.3%), which, according to (Wing & Kristofferson, 1973), points to a lack of variability in the timekeeper mechanism. These trials were retained in the analysis, but the corresponding clock delay estimates were set to zero. To examine the potential effects of these clock and motor delay adjustments, we conducted a parallel analysis in which we excluded the trials with serial correlations being positive or lower than −0.5. This showed that the adjustments had slightly increased the clock variance results (YA: 13.5%, OA: 6.5%, YA: 4,4%), however the proportional relationship between the groups remained largely consistent and did not affect the conclusion.

As a general sensorimotor measure, we analyzed the mean inter-tapping interval to evaluate the ability to maintain the target tapping speed.

### Grip force adjustment

In this task, we measured how participants modulated their grip force in function of changes in load force, when lifting a manipulandum. Grip force refers to the force applied to hold the object, while load force is the upward force required to lift the object. Two Nano 17 force sensors (ATI Industrial Automation) were mounted on the manipulandum. To enhance surface friction, sandpaper (P600 grit) was applied to the sensors. Forces were recorded along three axes at a sampling rate of 1000 Hz. The combined weight of the manipulandum and sensors was approximately 120 grams.

Participants were seated with the manipulandum positioned on a table in front of them. They were instructed to lift the manipulandum by gripping the force sensors between their thumb and index finger (Fig.1A). A set of stripes marked on a pole positioned on the table indicated the required height of lift for each trial. A sound beep indicated the start and end of a trial. The task comprised two conditions: the up-down condition and the hold condition.

**Figure 1.**
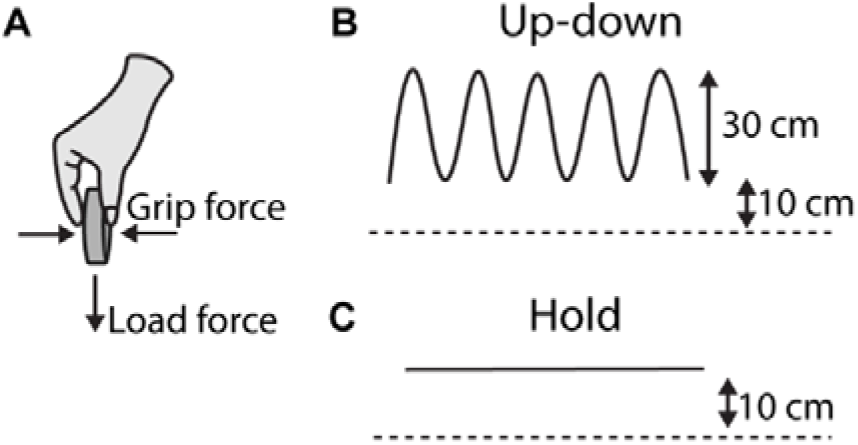
Grip force adjustment task. **(A)** Participants held the force sensor between the thumb and index finger. Grip force and load force were measured in X and Z directions, respectively. **(B)** In the up-down condition, the manipulandum was lifted 10 cm from the table (dashed line). From there is was moved up and down over a distance of 30 cm (black curves). **(C)** In the hold condition, participants held the manipulandum stationary at 10 cm above the table.

In the up-down condition, the participants cyclically moved the manipulandum up and down between two indicators with an in-between distance of 30 cm (Fig.1B). Movement speed was paced by a 2 Hz metronome. Each tone indicated either an upward or a downward movement, such that a full cycle consisted of two beats, corresponding to cyclic movements at 1 Hz. Each trial lasted 10 s, including approximately 10 up-down repetitions. 5 trial repetitions were performed.

During the hold condition, participants had to lift the manipulandum 10 cm from the table and hold it there for 10 s (Fig.1C). Given the greater stability of this condition, only 4 trial repetitions were conducted.

Grip force and load force signals were measured as the forces in the horizontal direction (X) and vertical direction (Z), respectively.

For the cerebellar-specific measure, the timing of anticipatory grip force adjustments for upcoming load force changes during the up-down movements were computed. Grip force and load force data from the up-down condition were preprocessed to reduce high-frequency noise by applying a fourth-order Butterworth low-pass filter to both, using Matlab’s butter function. A cut-off frequency of 2 Hz was chosen, to exclude noise caused from sources other than the frequency of the object movement. The filter coefficients were determined based on the chosen cutoff frequency and the sampling rate of the data (1000 Hz). The filter was applied in a zero-phase manner using MATLAB’s *filtfilt* function to avoid phase distortion. This two-way filtering process (forward and reverse) ensures the signal remains in phase, allowing accurate temporal alignment of force signals across trials.

Based on the filtered data, all load force maxima peaks within a 3 s to 9 s time window were identified, ensuring participants had started properly and excluding the last second of the original 10 s trial to capture approximately six full movement cycles per trial. Grip force maxima occurring within a 400 ms window centered around each load force peak (−200 ms to +200 ms) were designated as the corresponding grip force peaks. This window was chosen because, beyond approximately ±250 ms, the direction of the movement begins to reverse, making it unlikely that a true grip force peak associated with the current movement would occur outside this interval. Grip force maxima occurring exactly at ±200 ms from a load force peak were considered unreliable, as they were more likely to represent the highest value at the window boundary rather than a true peak, resulting in the removal of 342 trials (8.6%). To ensure data quality, participants were required to have at least 10 reliable grip force peaks remaining for inclusion in the analysis, which resulted in the exclusion of one participant. Average time differences between load force and corresponding grip force peaks were calculated and used as cerebellar-specific measures to evaluate grip force anticipation.

For the general sensorimotor measure, we calculated the grip force safety margins used by participants in the hold condition. Safety margins were defined as the amount of grip force exceeding the minimum required to hold the manipulandum, which was 1.18 N. We calculated this by subtracting the minimum required grip force value from the average recorded grip force overthe 3–6 s interval of each trial. Due to a procedural issue early in the study, where participants were allowed to contact the object before grip force baselines had stabilized, absolute grip force values are considered reliable only for a subset of participants: 14 young adults, 45 older adults, and 30 older-old adults. This issue was later corrected by adjusting the instructions to delay object contact until after proper baseline stabilization.

### Inter-joint coordination

In the inter-joint coordination task, anticipatory muscle activation was examined during arm movements, following the protocol of Maeda et al. (2017). Participants were seated in the Kinarm robotic exoskeleton (Kinarm, BKIN Technologies Ltd., Kingston, ON, Canada). The exoskeleton measures movement kinematics of the upper limbs. Additionally, surface electromyography (EMG) electrodes (Delsys system with Trigno Avanti Sensors) were placed on the skin to measure muscle activity of the biceps brachii long head (elbow flexor), triceps brachii lateral head (elbow extensor), pectoralis major clavicular head (shoulder flexor), and posterior deltoid (shoulder extensor) of the right arm.

Participants were seated in front of a screen and could not directly see their hand. Instead, a white cursor on the screen represented the hand’s position. They were instructed to move their right hand from a home position to a goal target using only elbow movements, while keeping their shoulder stable in an angle of 60° (Figure 2).

**Figure 2.**
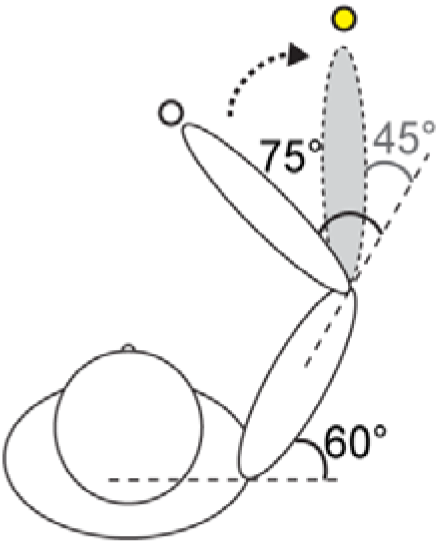
Inter-joint coordination task. The participant’s hand starts at the home position (white dot), with the elbow positioned at 75°. In this this example, the task requires an elbow extension movement towards the goal target on the right (yellow dot), reaching an elbow angle of 45°. Throughout the movement, the shoulder angle should remain fixed at 60°.

The home position was placed at an elbow angle of 75° (external elbow angle while shoulder remains at 60°). Two goal targets were used: a left target at 105° elbow angle and a right target at 45° elbow angle. This setup allowed participants to complete reaching movements with 30° pure elbow motion.

Four movement directions were tested: (1) from the home position to the left target (elbow flexion), (2) from the left target back to the home position (elbow extension), (3) from the home position to the right target (elbow extension), and (4) from the right target back to the home position (elbow flexion). This structure ensured that the goal target of each trial starting from the home position also served as the starting point for the subsequent return movement. The goal targets for trials initiating from the home position (directions 1 and 3) were presented in a pseudorandomized order.

At the start of each movement, the cursor disappeared and remained off until the trial ended, preventing feedback on hand position. Participants were instructed to stop their movement as close as possible to the goal target. Movement speed was targeted between 150–200 ms, with color-coded visual feedback indicating whether the speed was within the desired range (green), too slow (blue), or too fast (orange).

Prior to the inter-joint task, normalization trials were conducted to standardize muscle activity, with a value of 1 representing the mean activity required to counter a 2 N·m torque (Pruszynski et al., 2008)(Pruszynski et al., 2008). After becoming familiar with the target movement speed, participants completed 30 arm movement trials per condition, totaling 120 trials.

Trials in which movements started from the home position and proceeded to the left target (flexion) or right target (extension) were included in the analysis. The remaining trials were extensively analyzed in a separate project, taking into account the effect of the different home positions as well. Joint kinematics were sampled at 1000 Hz. EMG signals were sampled at 1000 Hz, bandpass filtered (20-100 Hz, 2-pass, 2nd-order Butterworth), full-wave rectified, and normalized so that a value of 1 corresponds to the mean activity of a given muscle sample during movements resisting a 2 Nm torque. The start and end of the elbow movement were defined as the moments when the elbow joint reached 5% of its peak angular velocity. As a cerebellar-specific measure, we analyzed the timing of anticipatory shoulder activation relative to the start of the movement, based on EMG measures. The first phasic EMG burst was scored based on the mean muscle activity of all trials in each movement direction. Baseline EMG activity was computed over a fixed time window of 100 ms, starting 400 ms before the start of the movement. The onset of the first phasic EMG burst was computed as the time at which the EMG signal rose 3 standard deviations above the mean baseline EMG level and remained above that level for at least 50ms (Maeda et al., 2017).

As a general sensorimotor measure, we analyzed deviations from the fixed 60° shoulder angle during elbow movement to assess mo vement stability and control. Total shoulder deviation was quantified as the absolute area between the deviation curve and the 60° reference line. Trials in which participants moved in the opposite direction of the intended movement were excluded from all analyses, which concerned 180 extension trials (3.75%) and 177 flexion trials (3.69%). Additionally, 44 participants (17 young, 24 older, 3 older-old) showed no EMG burst during extension trials, and 61 participants (18 young, 33 older, 11 older-old) showed no EMG burst during flexion trials, leading to their exclusion from the EMG analysis. In the result section we focused on the results from the extension movements (home position to right target) because of better EMG burst data quality compared to data from the flexion movements (which are described in the supplementary results). Detailed results of this task are describe in another paper (Matthijs et al., 2025)

### Eye movement coordination

The eye movement coordination task assessed eye velocity anticipation for a moving target, as well as the coordination between smooth pursuits and saccades during the visual tracking of that target while it was temporarily occluded, following the protocol of (Orban De Xivry et al., 2006).

For this task, we used the Eyelink 1000 Gaze-tracker installed on the Kinarm exoskeleton for target presentation and simultaneous gaze-tracking. Participants were seated in front of the screen, their foreheads resting against a support, with their gaze directed downward toward the display. Before task execution, a 5-point eye calibration was performed. During the task, participants were instructed to visually track a horizontally moving target on the display. The target was a red dot with a visual radius of 1 cm. Each trial began with the target appearing stationary at the starting position of its movement trajectory for 1 s. Just before the movement started, the target briefly disappeared for 0.3 s. The target always moved left to right in a straight line at a constant speed of 12 cm/s, covering a 28.8 cm in 2.4 s. After the movement ended, the target disappeared immediately, and participants’ gaze was tracked for an additional 0.3 s. After an inter-trial period of 1.5 s the next trial started (Figure 3).

**Figure 3.**
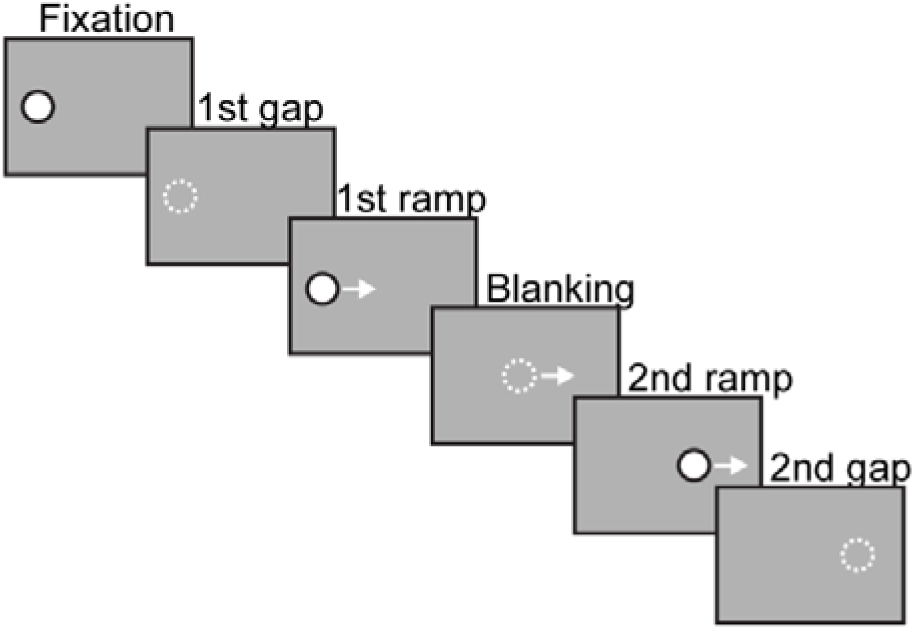
Eye movement coordination task. (1) The task starts with target fixation for 1 s. (2) followed by a brief disappearance of the target for 0.3 s. (3) When the target reappears it starts moving constantly from the left to the right side of the screen for 0.8 s. (4) In the blanking trials, the target is invisible for a part of the trajectory for 1 s. (5) For the last part of the moving trajectory the target reappears 0.6 s. (6) At the end of the trajectory the target disappears briefly before the next trial starts 0.3 s.

The task included two types of trials: control trials and blanking trials. During the control trials, the target was continuously visible after target motion onset (for 2.4 s). In blanking trials, the target remained visible for the first 0.8 s after movement onset, then disappeared for 1 s, and finally reappeared for the last 0.6 s of target motion. Participants were instructed to track the target as smoothly as possible during its visible phases and to continue tracking the target’s imaginary trajectory during the blanking periods, avoiding the tendency to make a rapid gaze shift to the expected reappearance location.

The experiment started with a baseline block of 20 control trials, designed to familiarize participants with the target trajectory. This was followed by five test blocks, each consisting of 26 trials: 6 control trials and 20 blanking trials. Each test always block began with two control trials, while the order for the remaining control and blanking trials were pseudorandomized. After completing each block, participants were allowed to rest for as long as needed before proceeding to the next block.

Eye-tracking data were collected at 1000 Hz. For analysis, the data was filtered with a low-pass, zero-phase autoregressive filter with a cut-off frequency of 20 Hz. Vectorial eye velocity was calculated based on only the horizontal displacement of the eye. To obtain smooth eye velocity, all saccades were removed along with 25 ms of data preceding and following each detected saccade and replaced by linear interpolation. Saccades were identified using an acceleration threshold of 3 cm/s² and a minimum duration of 30 ms.

Based on the control trials, in order to avoid potential effect of the upcoming blanking period, we computed anticipation of eye velocity for the upcoming target motion as a cerebellum-specific measure. Anticipatory eye velocity refers to the ability of the eyes to begin moving in advance of a predictable target, adjusting their velocity so that smooth pursuit is properly aligned with the target’s motion immediately upon its appearance or movement. This reflects the brain’s capacity to predict target motion and generate feedforward commands, reducing the lag that would otherwise occur if eye movements were driven solely by visual feedback.

Anticipatory eye velocity was calculated as the average smooth eye velocity between 50 and 100 ms after target motion onset. Considering the latency period of smooth pursuit (Rashbass, 1961), this interval captures the moment the eye velocity starts to increase in anticipation for the expected motion.

Based on the blanking trials, the synergy between saccades and smooth pursuits in the absence of visual guidance was computed as another cerebellar-specific measure. This synergy was defined as the regression sloop of smooth pursuit displacement to saccadic displacement during the blanking period, with a slope of −1 indicating perfect coordination.

We calculated each component separately, starting from 170 ms after occlusion onset (to account for typical saccade latency (Orban De Xivry et al., 2009) until the end of the blanking period. Saccadic displacement was computed as the sum of all saccadic amplitudes within this interval. Smooth pursuit displacement was obtained by integrating the smooth eye velocity signal over the same period. Both displacement measures were then normalized to the total eye displacement observed during the same time interval in a control condition.

We computed the visually guided eye velocity as a general sensorimotor measure, based on the control trials. For this, we calculated the mean smooth eye velocity between 800 and 850 ms after target motion onset. This interval captures the latest period of eye movements that is influenced by motion of the visual target, considering a delay of 100ms for a visual stimulus to influence the eye movement.

All trials underwent visual inspection to exclude those with blinks during critical periods of interest. These periods included the anticipation and movement phases for control trials, and the occlusion phase for blanking trials. On average, 92.00% (±11.51%) of control trials and 61.86% (±19.94%) of blanking trials (out of 50 and 100 trials, respectively) were included in the analysis for the young adult group. In older adults, 95.22% (±5.65%) of control trials and 66.50% (±19.81%) of blanking trials were included.

### Force matching

In the force matching task, we measured sensory attenuation during self-applied touch, based on the force-matching task protocol of Wolpe et al. (2016).

Following this protocol, a homemade set-up was built, which was placed on a table. It consisted of a 10 cm lever attached to a torque motor and connected to a tablet, which acted as a control panel. Participants sat in front of the force lever, with their left index finger placed underneath the lever with the pulp facing upwards. A force sensor at the end of the lever remained in contact with the finger, measuring the forces applied to the finger. White noise was delivered through headphones to mask any sounds from the torque motor that might help participants to estimate force output by ear instead of somatosensory perception.

Each trial began with a force perception phase which includes the application of a target force to the participant’s finger by the torque motor for 2.5 s. The target force was randomly chosen from four possible forces: 1 N, 1.5 N, 2 N, and 2.5 N, following the protocol of Wolpe et al. (2016). At the end of the force perception phase, a beep indicated the start of the force reproduction phase. During this phase, participants had to apply force to their own finger to match the perceived target force. There was no time limit for the reproduction phase. Participants said ‘yes’ when they felt the applied force matched the target force, prompting the instructor to press a button to mark the match. After this verbal signal, they were instructed to maintain the force for an additional 2.5 seconds until a second beep signaled that they could stop exerting the force.

There were three conditions for the reproduction phase: the slider condition, the button condition, and the lever condition. In the slider condition, participants controlled the torque motor with their right index finger using a slider (a linear potentiometer; Figure 4A). The slider condition served as a control because participants’ finger movement only indirectly determined the torque motor’s output, breaking the direct link between their own force production and the experienced force. This allowed us to compare responses to externally generated forces versus self-generated ones. In the button condition, participants pressed a button located above the lever with their right index finger (Figure 4B). The force exerted on the button was transmitted to the left finger via the torque motor, linking the force applied by the right index finger directly to the force applied on the left index finger (self-applied force). In the lever condition, participants pressed directly on top of the lever, mechanically transmitting the force to the left finger underneath the lever (self-applied force; Figure 4C).

**Figure 4.**
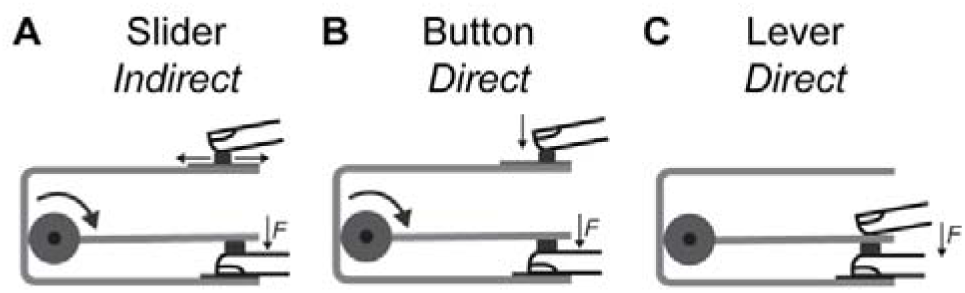
Force matching task. **(A)** In the slider condition (control), participants used the slider located on top of the device to manually control the force applied to the finger via the torque motor. **(B)** In the button condition, a button was used which directly applied the same amount of force to the finger via the torque motor as was applied to press the button. **(C)** In the lever condition, participants pressed the lever on top of the finger to apply the force.

All participants performed both the slider and button conditions. Interim data checks revealed that the young adult group did not use excessive self-applied force in the button condition, suggesting that they did not exhibit sensory attenuation. Consequently, the lever condition was added as an additional condition of self-applied force, to induce sensory attenuation effects in the young adults. The lever condition was performed by a subset of participants: 14 young adults, 45 older adults, and 29 older-old adults. The order of the conditions was counterbalanced across participants.

Before each condition started, participants underwent an initial familiarization phase of eight trials (two cycles of the four target forces). The main experiment comprised 32 trials (eight cycles) per condition, with a short break (±30 s) provided every two cycles. Each task cycle consisted of four trials, with one trial per target force presented in random order.

Both the target and the reproduced forces were measured by a sensor placed beneath the lever in contact with the participant’s index finger with a 1000 Hz sampling rate. The reproduced force was calculated as the average self-applied force measured within a 500 ms window centered on the match mark, spanning from 250 ms before, to account for the instructor’s button press delay, to 250 ms after, reducing the influence of force destabilization that can occur if the participant maintains the matched force for too long.

As a cerebellar-specific measure, we assessed sensory attenuation during self-applied force in the button and lever conditions. This was quantified as the difference between the target force and the force reproduced by the participant (matching error), with a reproduced force greater than the target force indicating sensory attenuation during self-applied touch. Similarly, the matching error for the slider condition was calculated and used as a reference value. Force overcompensation, reflecting the amount of sensory attenuation, in the button and lever conditions was normalized by subtracting the slider reference value from the matching errors of both conditions, to correct for individual baseline force overcompensation.

Sensory precision during the slider condition was computed as a general sensorimotor measure. This was calculated as the slope of the linear fit between target forces (X-axis) and reproduced forces (Y-axis), This slope reflects participants’ ability to discriminate between target forces and accurately scale the corresponding reproduced forces.

### Reach adaptation

In the reach adaptation test, implicit adaptation of reaching movements was assessed using the task-irrelevant clamped feedback protocol established by (Morehead et al., 2017).

Participants were seated in the Kinarm end-point robot (Kinarm, BKIN Technologies Ltd., Kingston, ON, Canada; Figure 2A). They were using the right arm for task execution. The position of the right handle was indicated by a cursor on the screen.

Participants were instructed to perform reaching movements with their right arm, starting from a home position and extending through a goal target. The home position was indicated by a yellow circle in the middle of the screen. Goal targets were located at four different positions around the home target along the four diagonal axes, each at a distance of 8 cm from the home target. To encourage movement consistency across participants, a point could be earned for correct movement speed, defined as a movement duration within a 200 to 350 ms time window.

During the baseline phase, participants smoothly moved their hand from the home target through the goal target, while the cursor indicated their hand position (Figure 5A). The baseline phase consisted of 40 trials (10 cycles of four trials, one per target direction). If the participant did not achieve the correct movement speed in more than 20 trials, the baseline session was repeated to stimulate familiarization with the expected movement speed.

**Figure 5.**
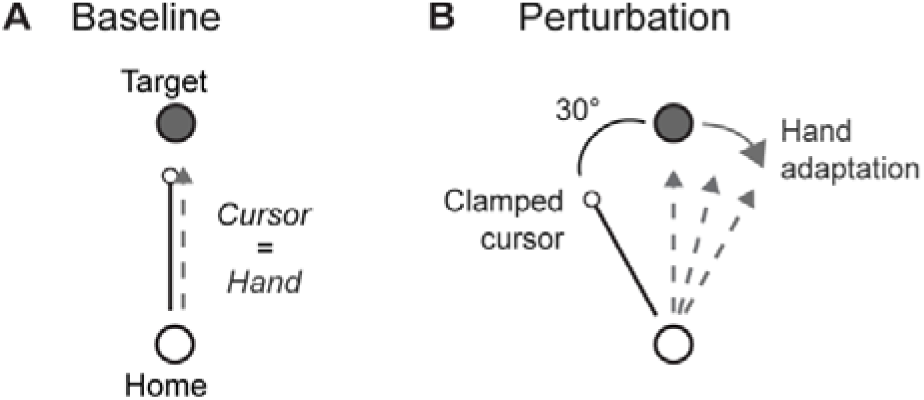
Reach adaptation task. **(A)** In the baseline condition, participants moved the cursor, representing their hand location, from the home position to the goal target. **(B)** In the perturbation condition, the cursor was artificially offset from the actual hand position, with a clamped deviation of 30° counterclockwise from the goal target. The dashed arrows indicate the adjustment of the hand in the opposite direction of the perturbation, demonstrating adaptive behavior to the perturbation over trials.

During the adaptation phase, the cursor’s angular direction was clamped 30° counterclockwise relative to the goal targets, while its radial position accurately reflected the hand position (Figure 5B). Participants were informed about the clamped cursor and instructed to move their invisible hand through the goal target, ignoring the cursor’s angular deviation but using it to gauge radial position. The adaptation phase consisted of 240 trials, with two 60-second breaks after trial numbers 120 (80 trials into the adaptation phase) and 200 (160 trials into the adaptation phase). The goal targets were presented in pseudorandom order during the baseline and adaptation phase.

All experimental conditions were programmed in MATLAB-Simulink (MathWorks, Natick, MA). Position data were sampled at 1,000 Hz. For each trial, the initial hand direction was calculated as the angle of the hand at 4 cm away from the home target as done in previous papers (Vandevoorde & Orban De Xivry, 2020). Hand error was determined by the difference between the angle of the initial hand direction and the angle of the goal target position. Trials with a hand error exceeding ±60° were excluded, as such large deviations typically reflected inaccurate reaching caused by predicting the target at an incorrect location, resulting in the removal of 618 trials (1.49%) from the dataset.

To establish a baseline reference, the average hand error for each participant was calculated from the last 20 baseline trials. Implicit adaptation to the perturbation was assessed as the change in hand error during the adaptation phase. The adaptation values were normalized by subtracting the baseline hand error from the adapted hand error to control for individual differences in initial reaching bias. This allowed us to isolate the change in performance specifically attributable to adaptation, rather than pre-existing error. Final hand adaptation was evaluated as the average adapted hand error across the last 40 perturbation trials and taken as a cerebellar-specific measure. The baseline hand error was taken as a general sensorimotor measure, reflecting reach accuracy.

### Speech adaptation

In the speech adaptation test, we investigated implicit speech adaptation in response to auditory perturbations using the abrupt perturbation protocol developed by (Parrell et al., 2017). This experiment tests whether and how individuals unconsciously adapt their pronunciation when the auditory feedback they receive through headphones is subtly altered to make their spoken verbs sound different.

The setup and software that were used for this task were similar to what is described in Parrell et al. (2017). Participants were seated in front of a computer screen that was connected to a laptop. They were wearing headphones that covered the ears. A separate microphone was positioned in front of the mouth. The head of the participant was resting against a head frame to minimize displacement between the microphone and the mouth. Speech was recorded with the microphone and processed by the computer.

Participants were instructed to read out loud the word that appeared on the screen in front of them. They received the auditory feedback of their own voice via the headphones, together with a slight noise, ensuring that the subjects heard themselves back only in the headphones and not directly. Each word was displayed on the screen for 1.5 s with an inter-trial interval of 0.75 s.

The experiment began with a familiarization phase of 21 trials, designed to test whether the auditory feedback was working properly and to help participants acclimate to the task. In each trial, one of the following words was randomly selected: ‘bid’ (b□d), ‘bed’ (b□t), or ‘bad’ (b□t). In Dutch, which was either the native language or a well-known second language for all participants, these are meaningful and commonly used words.

The familiarization phase was followed by a baseline phase that consisted of 30 trials with the word ‘bed’. During these trials, participants received accurate (veridical) auditory feedback about their own voice. During the adaptation phase, which lasted 90 trials, we altered auditory feedback by perturbing the first vowel formant (F1). Vowel formants are the resonant frequencies of the vocal tract that shape the acoustic quality of vowels. Each vowel has three formants, of which F1 is primarily related to vowel height with higher F1 values correspond to lower vowels, while lower F1 values correspond to higher vowels. We shifted F1 of the /□/ vowel by −125 Mels, which made the word ‘bed’ began to sound more like ‘bid’ (Figure 6).The perturbation was introduced abruptly at the start of the adaptation block, without informing participants. Lastly, a washout phase of 10 trials was included, during which veridical auditory feedback was restored. Throughout the task, participants were given short breaks (±5 s) every 25 trials, and they were instructed not to speak during these breaks.

**Figure 6.**
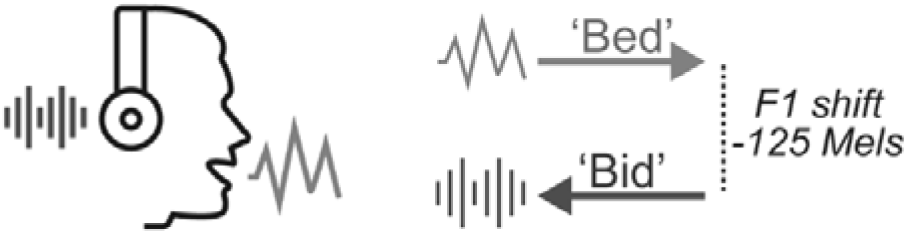
Speech adaptation task. During the baseline trials, participants pronounced the word ‘bed’ into a microphone and heard their own voice played back through headphones exactly as they had spoken it. During the perturbation trials, the first formant (F1) frequency of the pronounced word was altered, with a −125 Mel shift in the auditory feedback. As a result, when participants pronounced ‘bed’, they heard it as sounding more like ‘bid’.

The analysis software from Parrell et al. (2017) was used to extract the F1 values from the speech data (see supplementary methods I for detailed description). A baseline F1 frequency was established by calculating the median F1 value over the last 25 baseline trials. Instead of the mean, the median was calculated, because of the large trial-to-trial F1 variance in some subjects. For the same reason, we choose a larger baseline window compared to the study of Parrell et al. (2017) (baseline window of 10 trials), expecting the participants to be sufficiently familiarized with the task within the first five trials, since they were pronouncing a familiar word.

F1 implicit adaptation was assessed as the change in F1 during the perturbation phase. F1 adaptation values were normalized by subtracting the baseline reference F1 from the F1 adaptation values. Final F1 adaptation was calculated as the median of all F1 values across the last 30 adaptation trials and taken as a cerebellar-specific measure. As a general sensorimotor measure, we calculated the F1 variance during the baseline window, to assess speech stability.

### Mental rotation

In the mental rotation test, we tested the ability of participants to indicate whether a rotated letter was either normal or mirror-reflected using a protocol developed by McDougle et al. (2022), used with cerebellar patients. For this task, participants were sitting behind a table, using a laptop to perform the task.

In the task, the participants had to judge if a visual stimulus was presented normal (“R”) or mirror-reflected (“Я”). Eight capitalized sans-serif (Helvetica font) letter stimuli were used, consisting of normal and reflected versions of the letters F, G, J, and R. Letter stimuli were white and presented on a black background. Participants were instructed to press the left arrow key with their left index finger when the stimulus was in standard form and the right arrow key with their right index finger if the stimulus was mirror reflected (Figure 7). The stimulus was presented in the standard upright orientation (0°, baseline condition), or rotated at −135°, −75°, −15°, 15°, 75° or 135° and remained visible until one of the arrow keys was pressed. Stimuli were presented in a random order, with an equal number of normal and reflected presentations of each stimulus at each rotation sign and magnitude. After each trial, feedback on response accuracy was provided via on-screen text: a green ‘correct’ or a red ‘incorrect’ message displayed for 1 second. Prior to the start of the experimental block, the participants performed five practice trials to ensure that they understood the task instructions and were comfortably positioned to respond on the keyboard. The main task consisted of 144 trials, with an inter-trial interval of 2 s.

**Figure 7.**
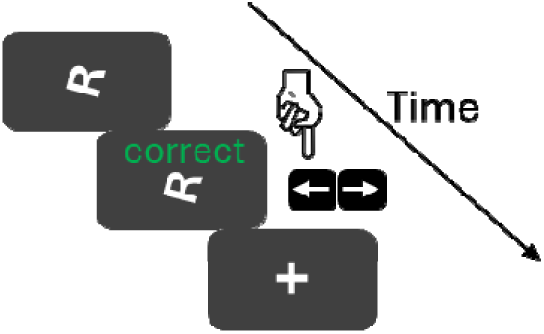
Mental rotation task. A rotated letter was presented on the screen that had to be judged whether it was presented normally or mirrored by pressing the left or right arrow button, respectively. Feedback about the correctness of the response was given afterwards, followed by a short fixation period before the next trial started.

Only data from the trials with correct responses were selected for the analysis, since errors in incorrect trials could not be clearly attributed to either difficulties with mental rotation or to impulsive key presses. For each of the rotations the mean reaction time was calculated, taken together the clockwise and counterclockwise (negative) angles as one rotation. A regression line was fitted through the reaction times at 0, 15°, 75° and 135°. The slope of this regression line reflects the rate of mental rotation, with steeper slopes indicating slower transformation per degree of rotation. Because the cerebellum is hypothetically involved in performing such continuous internal transformations, this slope was taken as a cerebellum-specific measure (McDougle et al., 2022; Picazio et al., 2013; Zacks, 2002). As a general sensorimotor measure, the baseline choice reaction time was computed, based on the mean reaction time in the absence of a perturbation (0°).

### MRI Acquisition

Images were acquired using a Philips 3T Achieva magnetic resonance scanner (Philips Healthcare, The Netherlands) with a 32-channel receiver head coil, located at the University Hospital of Leuven, Belgium. A high-resolution three-dimensional T1-weighted structural image was acquired (three-dimensional transient field echo (3DTFE); repetition time (TR) = 9.7 ms; echo time (TE) = 4.6 ms; inversion time (TI) = 900 ms; flip angle = 8°; voxel size = 0.89 × 0.89 × 1.0 mm; field of view (FOV)= 256 × 242 × 182 mm; 182 sagittal slices).

### Image processing

Before preprocessing, all scans were manually reoriented to the anterior commissure to optimize segmentation accuracy using SPM12 (Wellcome Trust Centre for Neuroimaging). An automated voxel-based morphometry (VBM) analysis was conducted using the Computational Anatomy Toolbox (CAT12.8.2) within SPM12. Structural T1-weighted images underwent brain tissue segmentation into gray matter, white matter, and cerebrospinal fluid, followed by spatial normalization to MNI space using the MN152 template. Total intracranial volume (TIV), total brain volume, and regional volumes by tissue type were extracted using the default CAT12 atlas-based parcellation. Total cerebellar gray and white matter volumes were computed by summing the respective cerebellar regions defined by the Computational Brain Anatomy (CoBrA) atlas (Park et al., 2014) which provides anatomically defined cerebellar parcellations.

Additionally, the interquartile range (IQR) metric, which assesses weighted image quality, was derived from the CAT12 analysis. Scans with IQR values below 4 were rated as good or excellent, which concerned all of the scans. To account for potential image quality differences between age groups, we statistically evaluated IQR values and included IQR as a covariate in the cerebellar volume analysis.

### Cam-CAN imaging data

In order to compare our cerebellar volume measurements with data from a different sample, we made use of structural T1-weighted images from the Cam-CAN repository (Shafto et al., 2014; Taylor et al., 2017). The images were acquired at the Medical Research Council (UK) Cognition and Brain Sciences Unit (MRC-CBSU) in Cambridge, UK, using a Siemens Magnetom TrioTim syngo MR B17 scanner (TR = 2250 ms, TE = 2.99 ms, TI = 900 ms, flip angle = 9°, voxel size = 1.0 × 1.0 × 1.0 mm, FOV = 256 × 240 × 192 mm, multi-slice mode). The dataset included images from 653 participants aged between 18 and 88 years, with approximately 100 individuals per decade. Voxel-based analysis was conducted and cerebellar gray and white matter volumes were computed using the same approach as described above, although performed separately from the images we obtained. To examine gray and white matter volumes and their relationship with age, which is expected to follow a quadratic pattern (Hoogendam et al., 2012; Walhovd et al., 2011), we used MATLAB’s *polyfit* function with a polynomial degree of 2. This function fits a second-degree polynomial to the data using a least-squares approach, generating a quadratic regression line to model nonlinear trends in volume changes with age.

### Statistical analyses

All statistical analyses were conducted using MATLAB R2022b (MathWorks, Natick, MA). To account for variability across age groups and minimize the impact of outliers in the behavioral data, robust statistics were employed using MATLAB’s *fitlm* function to fit a robust linear regression model:

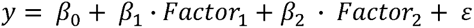

Age groups were included as two categorical predictors, coded using contrast coding: Factor 1 was coded as −1 (young), 0 (older), and +1 (older-old), capturing the linear trend across age groups. Factor 2 was coded as −1 (young), +1 (older), and 0 (older-old), enabling a direct comparison between the young and older groups, independent of the older-old group. This two-factor coding scheme also allowed for a separate post hoc comparison between the older and older-old groups via the appropriate linear combinations of model coefficients. In the model, *β*_0_ represents the intercept, *β*_1_ and *β*_2_ capture the effects of the age group factors 1 and 2 on the dependent variable, and ε represents the residual error term. The default settings for robust regression were applied, using an iteratively reweighted least squares method that assigns a weight to each data point. This weighting is incorporated as a scaling factor during the fitting process, improving the accuracy of the fit. Low-quality data points, such as outliers, receive lower weights, thus reducing their influence on the final parameter estimates. Overall behavioral and structural age group effect estimates are based on the statistical outcomes of the general linear model.

Two-sided post-hoc tests were conducted for each measure, including instances without a significant main effect of age, to quantify effect sizes that are used to provide a comprehensive overview of all results. Estimates from post-hoc testing between age groups were based on the estimated coefficient results of the linear model. Significance thresholds of the corresponding p-values were corrected for multiple comparisons using the False Discovery Rate correction. For effect sizes of post-hoc comparisons, the robust Cohen’s d was calculated based on t-statistics (t) of between-group effects, calculated as: 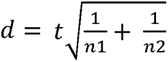, in which n is the group size. Corresponding confidence intervals were based on the standard error of d, using 95% confidence.

One-sided Bayes Factor (BF) based on a Cauchy prior with a width of 0.707 were computed thanks to the JASP software (version 0.19.3). To match the robust analyses performed above, we first winsorized the data (cutoff: 20%) in order to reduce the possible influence of outlier data. As in the JASP program, we interpret Bayes Factors (BF) as follows: a BF below 0.1 indicates strong evidence that the data from the older group do not differ from those of the young adults; a BF below 0.333 suggests moderate evidence for no difference. Conversely, a BF above 3 indicates moderate evidence for a difference between the groups, and a BF above 10 constitutes strong evidence for such a difference. Bayes factors between 0.333 and 1 or between 1 and 3 only provide anecdotal evidence in favor of no difference or difference, respectively, between the two age groups.

Additionally, we estimated split-half reliability for each behavioral measure to index within-participant consistency across trials. The grip force safety margin measure was excluded because it was derived from enduring grip force estimates based on only four trials. For each remaining task, trials were divided into odd and even subsets, outcome measures were averaged within each subset, and the correlation between odd- and even-trial averages was computed. Reliability estimates were corrected using the Spearman–Brown formula.

### Data availability statement

The data that support the findings of this study are not currently openly available due to the ongoing additional analyses/publications. They will be made available upon completion of the PhD theses of the first two authors. Data supporting the statistical analyses are available from the corresponding author upon reasonable request. Data are located in controlled access data storage at KU Leuven.

## Results

In this experiment, we tested a set of motor and cognitive tasks on a large group of young adults, older adults between 55 and 70 years old, and older-old adults above 80 years old. This test battery spans the diversity of functions attributed to the cerebellum. The tasks were selected as, for each of them, we could isolate one or several outcomes that can be linked to cerebellar functions because of their impairment in patients with cerebellar degeneration, or because of theoretical interpretations of the task outcome based on previous studies. Next to these cerebellar-specific outcomes, we also report general sensorimotor measures of motor functions for the same tasks.

**Figure 8.**
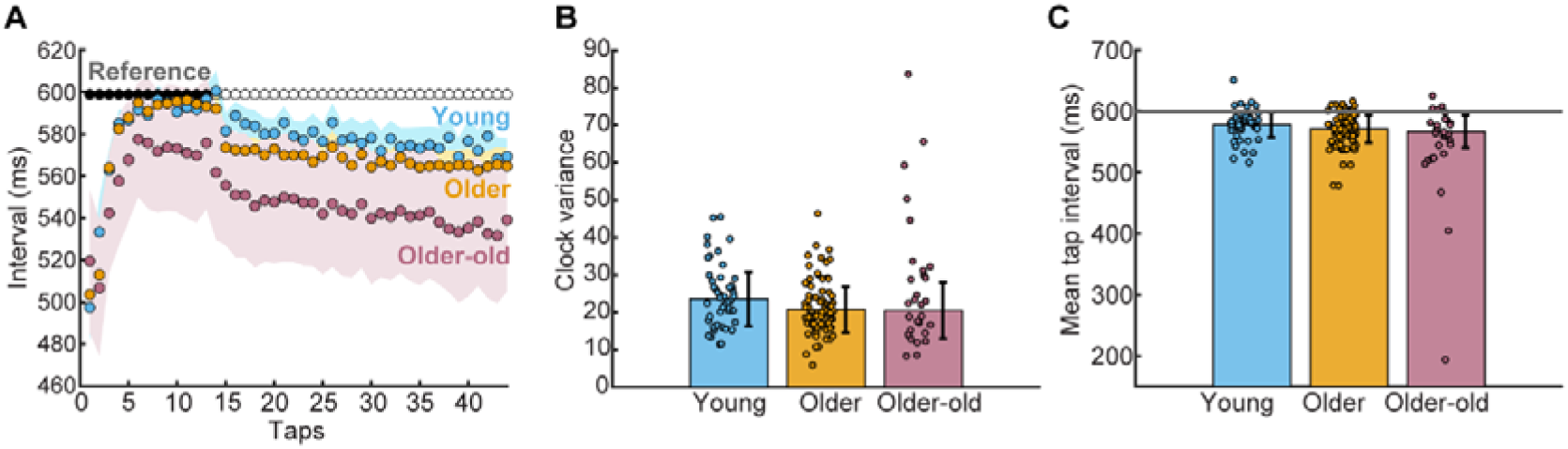
Rhythmic finger tapping results. **(A)** The average finger tap intervals per group, with shaded areas indicating 95% confidence intervals. The black dots represent the reference tapping interval (600 ms) from the initial auditory rhythm, which was played for the first 13 tones before becoming silent. Empty dots indicate the timing of the remaining, silent beats. **(B)** The clock variance, based on the last 30 unguided taps. **(C)** The mean tap intervals, based on the last 30 unguided taps shown with the reference interval (gray line). Bar heights and error bars represent robust means and standard deviations, respectively. Dots show individual participant scores. Analysis based on the data from 48 young adults, 77 older adults and 29 older-old adults.

### Rhythmic finger tapping

The timing function of the cerebellum, an important basis for the generation of accurate movements, was evaluated using a rhythmic finger-tapping task. Participants were presented with an auditory rhythm and instructed to tap in synchrony with it. After the sound stopped, they were required to continue tapping while maintaining the reference rhythm as accurately as possible.

As shown in **Error! Reference source not found.**A, participants needed about five taps to synchronize with the sound. Clock variance values were calculated based on the inter-tapping intervals of the unguided taps and taken as a cerebellar-specific measure, as it reflects the variability in the internal timekeeping mechanism.

Clock variances differed across age groups (Main effect of age: F(2,151) = 6.24, p = 0.002), with older adults showing lower variance (20.79 ± 6.11) compared to young adults (23.60 ± 7.21). All behavioral outcome measures, including those reported here, were analyzed using robust linear regression models to reduce influence of outliers. Post-hoc testing, based on the estimated coefficients of the linear model, showed this difference was insignificant (t(123) = 1.90, p = 0.06, d = 0.35; **Error! Reference source not found.**B). Clock variance for older-old adults (20.48 ± 7.51) was similar to older adults (t(104) = 0.17), p = 0.86, d = 0.04, BF = 0.052). The difference between older-old and young adults did not reach significance (*t*(75) = 1.64, *p* = 0.11, *d* = 0.39), although the older-old group showed numerically lower variance. Given the larger variance in the older-old adults, the evidence for no effect was, following the Bayesian analysis, only anecdotal (BF = 0.766). These results suggest that the timekeeping mechanism does not decline with age, not even in older-old adults.

As a general sensorimotor measure the average duration of the tapping intervals was computed based on the unguided taps. This reflects overall rhythmic finger-tapping performance, encompassing neocortical sensorimotor functions like finger movement (De Guio et al., 2012). The average duration of the tapping intervals differed across age-groups (F(2,151) = 4.57, p = 0.01; **Error! Reference source not found.**C). The young adults deviated the least from the reference rhythm of 600 ms (577.72 ± 20.57 ms). The older adults were slightly faster (570.73 ± 22.63 ms; t(123) = 1.37, p = 0.17, d = 0.25, BF = 5.239), followed by the older-old adults (567.00 ± 26.43 ms; vs. young: t(75) = 1.64, p = 0.10, d = 0.39, BF = 65.9); vs. older: t(104) = 0.62, p = 0.54, d = 0.13). However, post-hoc comparisons between age groups were not significant, indicating that the overall effect may reflect subtle differences not detectable in individual group contrasts.

These results suggest that sensorimotor tempo control is broadly preserved with age, though young adults may be marginally closer to the reference rhythm.

### Grip force adjustment

In the grip force adjustment task, we examined how anticipatory grip force adjustments are implicitly timed in relation to arm movements in the up-down direction. In this task, participants held a grip force sensor between their thumb and index finger while performing repetitive vertical arm movements or by lifting the object and holding it stationary. The timing of grip force anticipation relative to upcoming load force changes was assessed as a cerebellar-specific measure, by calculating the time difference between the peaks of grip and load force (Fig.9A). Accurate anticipation of grip force depends on cerebellar predictions of the load forces associated with self-generated movements (Kawato et al., 2003; Rost et al., 2005).

**Figure 9.**
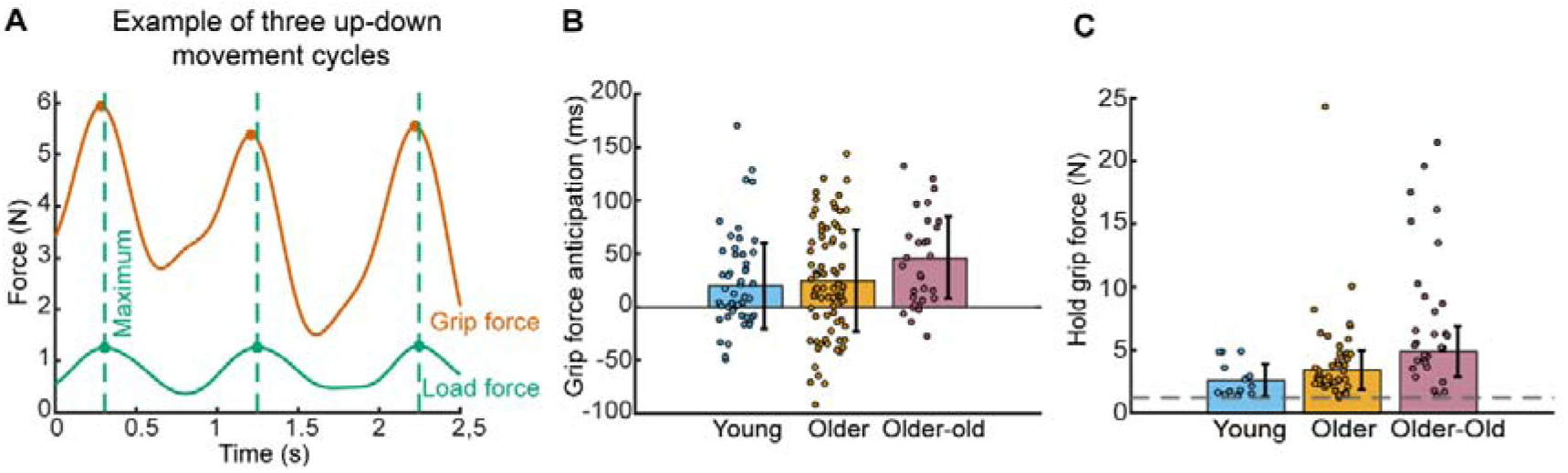
Grip force adjustment results. **(A)** Illustration of grip force and load force peaks during three up-down arm movements. Green dots mark load force maxima, with green dashed lines indicating their timing. Orange dots represent grip force maxima, occurring just before the load force peaks. **(B)** Grip force anticipation is the time difference between grip and load force peaks; positive values indicate grip force peaks occurred earlier (i.e., anticipation). Analysis based on the data from 49 young adults, 77 older adults and 30 older-old adults. **(C)** Average grip forces applied during the hold condition, displayed with the minimum required grip force indicated (dashed gray line). The extent to which the applied grip force exceeds this minimum reflects the grip force safety margin. Analysis based on the data from 14 young adults, 45 older adults and 30 older-old adults. Bar heights and error bars represent robust means and standard deviations, respectively. Dots show individual participant scores.

Across trials, grip force peaks consistently preceded load force peaks in all age groups, demonstrating the presence of anticipatory grip force adjustments to predicted load changes (positive bar heights in Fig.9B). No difference in the temporal advance in grip force anticipation was found across age groups(F(2,153) = 1.76, p = 0.18). Although it seemed that older (23.25 ± 47.69 ms) and older-old adults (40.60 ± 40.73 ms) tended to anticipate the load force peak more than young adults (20.81 ± 39.30ms), the Bayesian analysis provided moderate to strong evidence for an absence of difference in grip force anticipation (young vs. older: BF = 0.153, young vs older-old: BF = 0.095)

Excessive grip force, reflecting a safety margin, was analyzed during hold condition trials as a general sensorimotor measure. Although cerebellar patients also exhibited larger safety margins, these are thought to reflect a compensatory response to reduced sensorimotor performance rather than a primary cerebellar deficit, as similar compensatory increases are observed in motor deficits following cerebral damage (Hermsdörfer et al., 2003; Rost et al., 2005). The minimum required grip force to hold the object was 1.18 N, therefore any additional force corresponds to the safety margin used by the participants. This safety margin varied across age groups (main effect of group: F(2,86) = 19.6, p < 0.001; Fig.9C). Young adults exhibited the lowest grip forces (mean grip force: 2.73 ± 1.30 N; safety margin: 2.55 N). In contrast, older adults showed increased grip forces (mean: 4.57 ± 1.57 N; safety margin: 3.39 N), though this difference was not statistically significant, likely due to the small sample size of 14 young adults (t(57) = 1.2, p = 0.23, d = 0.52, BF = 3.8). The older-old adults displayed the highest grip forces (mean: 6.05 ± 2.02 N; safety margin: 4.87 N), which were significantly greater than those of young adults (t(42) = 3.15, p = 0.002, d = 1.10, BF = 56.26) and older adults (t(73) = 2.77, p = 0.007, d = 0.86). These results show that older and older-old adults maintain anticipatory grip adjustments but apply larger safety margins.

### Inter-joint coordination

In the inter-joint coordination task, we examined the role of timing mechanisms based on cerebellar predictions during pure elbow movements. Here, we looked at the timing of the shoulder muscle activation that is needed to compensate for the interaction torques, and to keep the shoulder stable.

Across all age groups, a consistent muscle activation pattern emerged: the shoulder antagonist activated first, followed by the elbow agonist, and then the movement was initiated (Fig.10A). The amount of shoulder anticipation was quantified as the time interval by which shoulder muscle activation onset preceded elbow muscle activation onset, and taken as a cerebellar-specific measure, as it reflects the cerebellum’s critical role in coordinating predictive motor timing and feedforward control of intersegmental dynamics (Bastian et al., 1996). No difference in shoulder muscle anticipation timing was found between age groups F(2,113) = 0.54, p = 0.59; Fig.10B). Shoulder muscle anticipation timing in young adults (13.67 ± 17.62 ms) was comparable to that in older adults (13.70 ± 19.66 ms), with the older-old adults showing slightly earlier anticipation (18.01 ± 19.64 ms), although not statistically meaningful. This suggests that shoulder muscle anticipation preceding elbow muscle activation remains intact in older and older-old adults (young vs. older: BF = 0.225; young vs. older-old: BF = 0.154).

**Figure 10.**
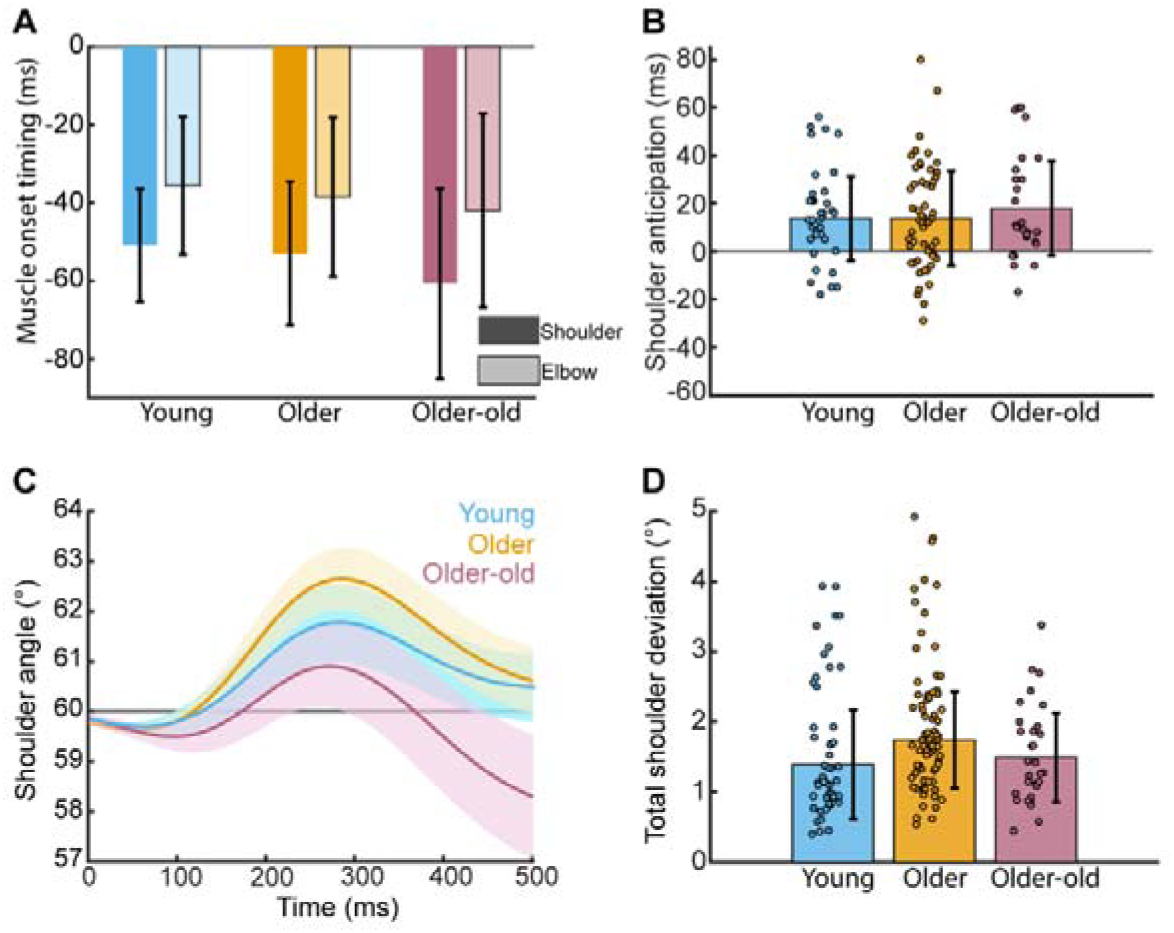
Inter-joint coordination results. **(A)** The timing of the activation of shoulder and elbow muscles before movement onset (at 0 s). Shoulder onset timing is indicated by the darkest color; elbow onset timing is indicated by the lighter color. **(B)** Shoulder anticipation timing indicates the time interval between the onset of shoulder muscle activity and the subsequent onset of elbow muscle activity. Analysis based on the data from 33 young adults, 56 older adults and 27 older-old adults. **(C)** During elbow movement execution, the shoulder deviated from the fixed 60° angle (gray line) across all three age groups. **(D)** Total shoulder deviation during the movement. Analysis based on the data from 50 young adults, 80 older adults and 30 older-old adults. Bar heights and error bars represent robust means and standard deviations, respectively. Dots show individual participant scores.

As a general sensorimotor measure we calculated the total amount of unintentional shoulder deviation from the fixed angle (60◦) during elbow movement, to assess joint stability and motor coordination (Fig.10C). Greater shoulder deviation indicates less efficient movement, as pure elbow motion would have been sufficient to reach the target. Such deviation may arise from factors like mobility impairments that commonly accompany aging (Davis, 2023). A main effect of age on shoulder deviation was found (F(2,157) = 5.96, p = 0.003; **Error! Reference source not found.**D). Young participants showed the least deviation from the fixed shoulder angle (1.39° ± 0.78°), with slightly greater deviation observed in older adults (1.74° ± 0.69°; t(128) = 2.35, p = 0.02, d = 0.42, BF = 3.7). However, this difference did not remain significant after correction for multiple comparisons. Total shoulder deviation in older-old adults (1.48° ± 0.64°) was not different from either young adults (t(78) = 0.53, p = 0.60, d = 0.12, BF = 0.229) or older adults (t(108) = 1.41, p = 0.16, d = 0.30). However, the pattern of deviation in the older-old adults was different (**Error! Reference source not found.**D), showing more deviation against the direction of the movement (<60°), while young and older adults showed more deviation in the direction of the movement (>60°). Note that results from the flexion movement data support our findings of preserved shoulder anticipation and increased shoulder deviation in the older age groups (see supplementary results).

The inter-joint results showed that predictive motor control, as reflected by preserved shoulder muscle anticipation was still intact in old and older-old adults. In contrast, age-related differences emerged in joint stability, with older adults showing slightly increased unintentional shoulder deviation during elbow movement, indicative of subtle changes in sensorimotor coordination, despite preserved anticipatory control.

### Eye movement coordination

Beyond anticipation involved in inter-joint coordination, we also investigated anticipatory eye velocity for a predicted moving target during the eye movement coordination task. The participants visually tracked a target moving at a constant speed from the left to the right side of the screen. Prior to movement, the target appeared at the starting position before briefly disappearing and reappearing at target motion onset. During this gap period, participant’s eye velocity increased in anticipation of upcoming target motion onset (Figure 11A). This anticipation started approximately 0.2 s before target motion onset in both young and old adults (older-old adults were not tested, see methods). The average velocity reached at the end of the gap period (and thus, at the time of target motion onset), was calculated as a cerebellar-specific measure, as this anticipatory eye velocity was previously found to be reduced in cerebellar patients compared to healthy control (Lekwuwa et al., 1995; Moschner et al., 1999). There was no difference in anticipatory eye velocity between young (3.54 ± 0.91cm/s) and older adults (3.63 ± 0.97cm/s; t(120) = 0.44, p = 0.66, d = 0.08; Figure 11B). This provides moderate evidence that the ability to anticipate for upcoming target motion was intact in older adults ( BF = 0.125).

**Figure 11.**
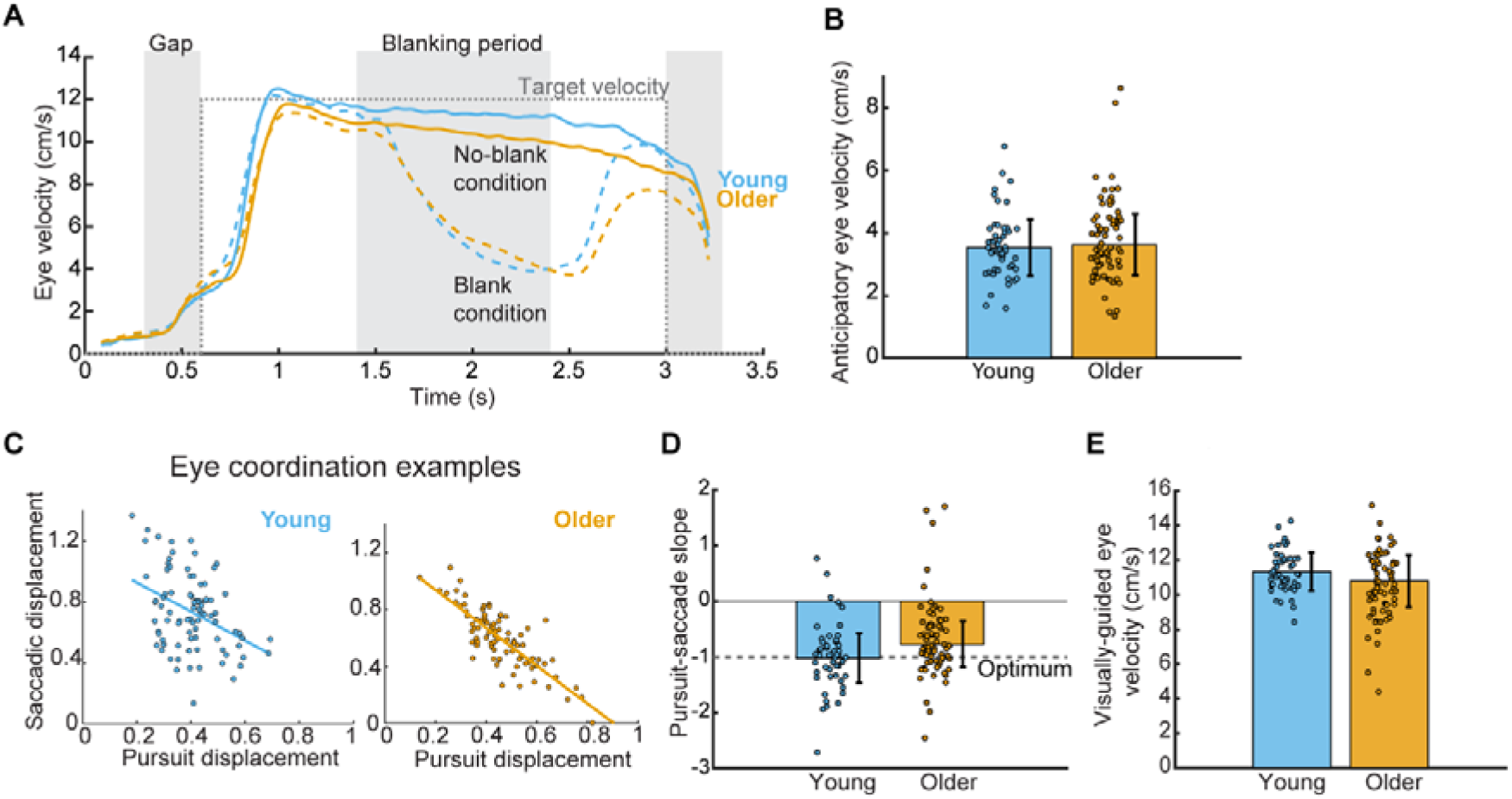
Eye movement coordination results. **(A)** The average eye movement velocity over trials for young and older adults. The dashed gray line represents the target speed, showing that the target remains stationary initially, then starts moving, and finally comes to a stop again. The gray areas indicate phases where the target was invisible, with the middle gray block representing the blanking period during which the target disappeared while moving (blank condition). Eye velocity in the blank condition (dashed colored lines) was decreased in both age groups compared to eye velocity during no-blank control trials (solid-colored lines), when the target remained visible for the whole trajectory. **(B)** Anticipatory eye velocity for the expected upcoming target motion. Analysis based on the data from 49 young adults, and 73 older adults. **(C)** Two examples of a young (left) and older participant (right) of saccadic and smooth pursuit displacements relative to total target displacement across individual trials (dots), during the blanking period. The fitted line represents the synergy between saccades and smooth pursuits, indicating mutual compensation. **(D)** Synergy between smooth pursuits and saccades during the blanking period. Analysis based on the data from 49 young adults, and 72 older adults. **(E)** Visually-guided eye velocity during eye tracking in the no-blank (control) condition. Analysis based on the data from 49 young adults, and 73 older adults. Bar heights and error bars represent robust means and standard deviations, respectively. Dots show individual participant scores.

In the same task, we examined the coordination of saccades and smooth pursuit as a second cerebellar-specific measure, as the generation of predictive saccades involves cerebellar computations (O□Driscoll et al., 2000). However, there is still no definitive evidence of deficits in cerebellar patients. During the blanking period (i.e. when the moving target was invisible), smooth eye velocity decreased (Figure 11A) and saccades occurred. Despite the lack of visual input, the amplitudes of the saccades and the decrease in eye velocity were negatively correlated, indicating coordination between the two types of eye movements based on extraretinal signals. The amount of the coordination is reflected in the negative regression coefficient between smooth pursuit and saccades (Figure 11C).

A difference in regression coefficients was found between the two age groups (t(119) = 2.71, p = 0.008, d = 0.5, BF = 106.8), with young adults (mean: −1.02 ± 0.44), being closest to the optimal value of −1, indicating highly effective coordination between the two types of eye movements (Figure 11D). In contrast, older adults exhibited a shallower slope (−0.77 ± 0.41), suggesting reduced coordination efficiency between the two types of eye movements. For the no-blanking control condition, we defined visually-guided smooth pursuit performance as a general sensorimotor measure (Figure 11A, solid lines). Visually guided smooth pursuit relies on a cortical initiation network with projections to the basal ganglia and cerebellum (Frei, 2021; Thier & Ilg, 2005). Participants from both age groups demonstrated eye velocities approaching the target speed of 12 cm/s. While the age effect did not reach statistical significance (t(120) = −1.86, p = 0.07, d = 0.34, BF = 12.647), young adults exhibited a slightly higher pursuit gain, with average eye velocity closer to the target speed (11.33 ± 1.11 cm/s) compared to older adults (10.80 ± 1.51 cm/s; Figure 11E). This trend may indicate a subtle age-related reduction in visually guided pursuit performance. Together, these findings suggest that while anticipatory eye movements and visually guided pursuit are largely preserved in older adults, age-related differences emerge in the coordination between saccades and smooth pursuit, indicating a specific decline in predictive sensorimotor integration.

### Force matching

Just as successful visual tracking relies on the coordination between saccades and smooth pursuit, force production depends on the interaction between sensory input processing and predictions about expected motor output. During self-applied touch, the perceived pressure results from subtracting cerebellar predictions, based on an efference copy of the motor command, from afferent somatosensory signals (Cao et al., 2017; Kilteni et al., 2020; Wolpe et al., 2016). Because of this predictive subtraction, a greater amount of force must be applied to perceive the same level of pressure during self-touch compared to touch originating from an external source.

Participants were able to accurately match an external force by manipulating a slider with their right hand, which controlled a torque motor applying force to the left finger via a lever (slider condition). In this control condition, young adults were the most accurate (mean reproduction error: −0.05 ± 0.22 N, dashed blue line in Fig.12A). In contrast, older and older-old adults slightly underestimated the force when using the slider (older: −0.24 ± 0.24 N; older-old: −0.27 ± 0.27 N; orange and pink dashed lines in Fig.12A).

In contrast, participants exerted more force when they controlled the lever-applied force by pressing on a force sensor positioned directly above the lever, using their right hand. The self-applied force was perceived as weaker, leading participants to apply more force than required on the force sensor (direct condition; solid lines in Fig.12A). This force overcompensation, taken as a cerebellar-specific measure, differed between age groups (F(2,154) = 28.05, *p <* 0.001; Fig.12B) after normalization for performance in the slider condition. We found little to no overcompensation in young adults (−0.01 ± 0.28 N). In contrast, overcompensation was significantly greater in older adults (0.31 ± 0.31 N; t(126) = 5.14, p < 0.001, d = 0.94, BF = 0.028), and even more pronounced in older-old adults (vs. young: t(75) = 7.22, p < 0.001, d = 1.70, BF = 0.133; vs. older: t(107) = 3.50, p < 0.001, d = 0.76). This shows older and older-old adults exhibited more force overcompensation, suggesting they experienced larger sensory attenuation than the young adults. A third condition tested on a subset of participants demonstrate that this paradigm can produce force overcompensation in the young adults as well (see supplementary results on the lever condition).

**Figure 12.**
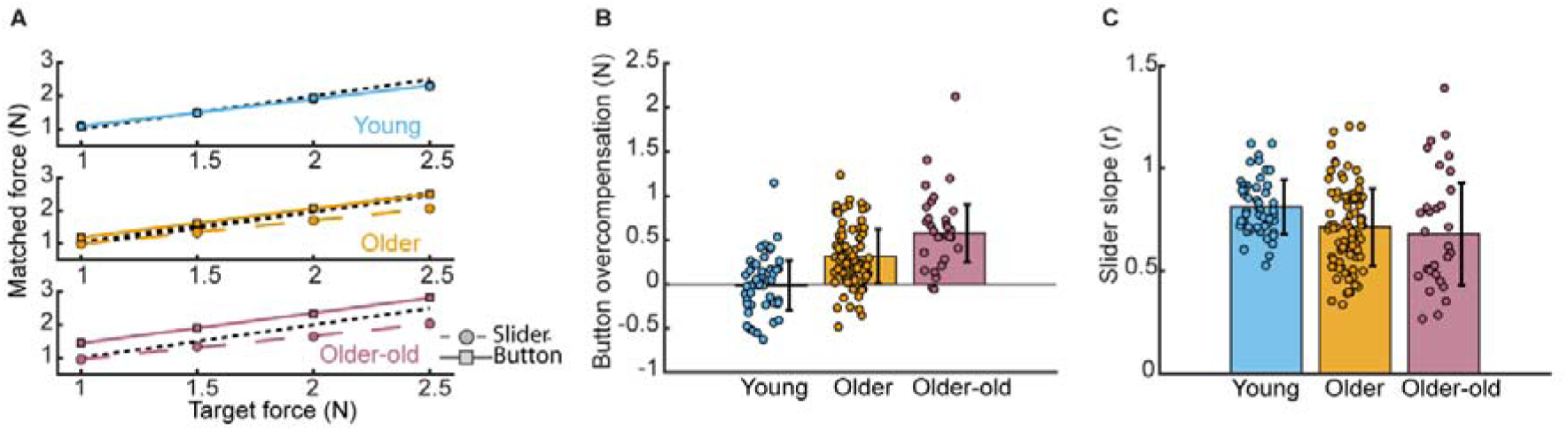
Results from the force matching test. **(A)** Mean linear regression fits of the matched forces against each of the target forces for each age group. The dashed black line is a reference to indicate perfect correspondence between matched and target forces. The dashed colored line with corresponding circles, show the results in the slider condition, which was used as a reference. The solid-colored line and corresponding squares show the button condition results. **(B)** Average overcompensation in the button condition. **(C)** Average slopes of linear relationship between target forces and matched forces during the slider condition, indication sensory sensitivity. Bar heights and error bars represent robust means and standard deviations, respectively. Dots show individual participant scores. Analysis based on the data from 48 young adults, 80 older adults and 29 older-old adults.

Greater overcompensation in older and older-old adults may stem from increased reliance on cerebellar predictions due to noisier somatosensory signals (Wolpe et al., 2016). To assess sensory sensitivity, we analyzed the slope of the relationship between target forces and reproduced forces in the slider condition and took it as a general sensorimotor measure. Shallower slopes indicate reduced sensitivity to force differences, which likely reflects sensory rather than cerebellar contributions (Wolpe et al., 2016). Our analysis revealed a significant effect of age on force sensitivity (F(2,154) = 4.8, p = 0.009). Young adults exhibited the steepest slopes, being closest to 1 (0.81 ± 0.13; Fig.12C). Compared to the young adults, older adults demonstrated significantly shallower slopes (0.71 ± 0.19; t(126) = −2.58, p = 0.01, d = 0.04, BF = 91.3), being similar to the slope of the older-old adults (0.68 ± 0.25; vs. young: (t(75) = −2.67, p = 0.008, d = 0.09, BF = 17.3; vs. older: t(107) = −0.72, p = 0.47, d = 0.06). Although the slopes in older and older-old adults were significantly shallower compared to young adults, the small effect sizes reveal that this is only a subtle difference.

### Reach adaptation

Prediction about the sensory consequences of one’s own movement also plays important role in motor adaptation as well (Morehead et al., 2017). When a task-irrelevant clamped feedback perturbation was introduced, participants from all age groups gradually adapted their reach direction (Figure 13A). In the final adaptation phase (last 40 trials), taken as a cerebellar-specific measure, older and older-old adults appeared to exhibit larger adaptation (21.08 ± 7.09° and 21.47 ± 7.62°, respectively) than younger adults (18.64 ± 7.50°), but this difference was not confirmed statistically (F(2,157) = 5.52, *p* = 0.08; Figure 13B). These findings suggest that implicit motor adaptation remains intact with advancing age (young vs older: BF = 0.079, young vs older-old: BF = 0.105).

**Figure 13.**
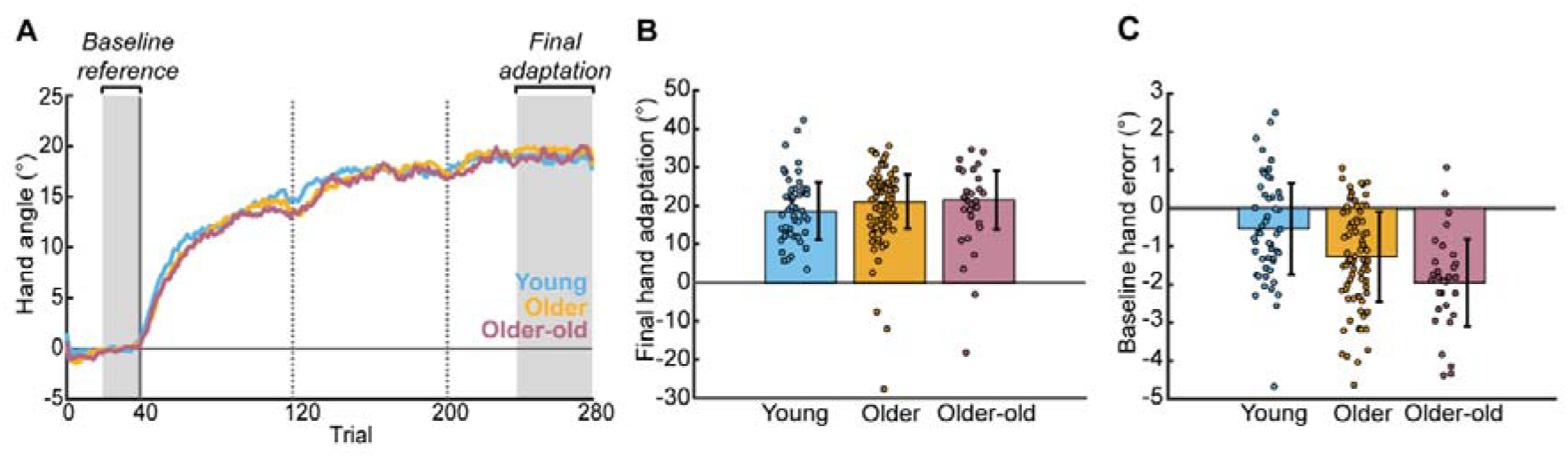
Results from the reaching adaptation task. **(A)** Trajectory of average baseline-corrected hand adaptation across all trials, per age group. Baseline correction used the final 20 trials of the baseline phase as a reference (shown by the first gray area). Perturbation began after trial 40 (vertical solid line). Breaks were provided at 80 and 160 trials after the onset of perturbation (dashed vertical lines at trials 120 and 200, respectively). The average late hand adaptation was calculated from the final 40 perturbation trials, referred to as the late adaptation period (second gray area). **(B)** Final hand adaptation. **(C)** Reach accuracy during the baseline phase. Bar heights and error bars represent robust means and standard deviations, respectively. Dots show individual participant scores. Analysis based on the data from 50 young adults, 80 older adults and 30 older-old adults.

Reaching accuracy during baseline trials, used as a general sensorimotor measure, showed that all age groups were able to reach near the target though significant differences in accuracy were observed between age groups (F(2,157) = 10.8, p < 0.001). Young adults demonstrated the highest accuracy (mean hand error: −0.55 ± 1.20°), showing a lower mean error than older adults (−1.27 ± 1.18°; t(128) = 2.99, p = 0.003, d = 0.54, BF = 152.3; Figure 13C). Older-old adults were the least accurate (−1.96 ± 1.14°), exhibiting significantly greater error than both young adults (t(78) = 4.54, p < 0.001, d = 1.05, BF > 1000) and older adults (t(108) = 2.39, p = 0.02, d = 0.51). Together, these results suggest that while aging is associated with a modest decline in baseline reaching accuracy, implicit motor adaptation remains well preserved in older and older-old adults.

### Speech adaptation

Next to reach adaptation, we investigated cerebellar-dependent speech adaptation by introducing a perturbation to the first formant (F1) of vowel sounds while participants spoke the word “bed” (Parrell et al., 2017). Due to the perturbed auditory feedback, they heard themselves saying “bid,” prompting adaptive changes in their speech production (Figure 14A).

**Figure 14.**
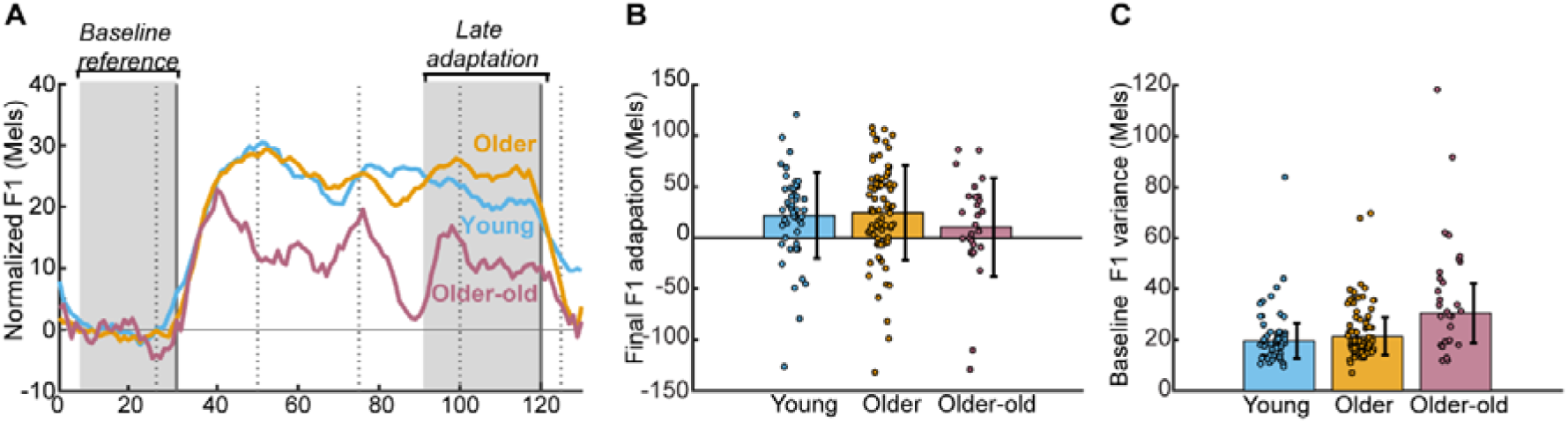
Results from the speech adaptation task. **(A)** Trajectory of the baseline-corrected F1 trajectories. Baseline correction used the final 30 trials of the baseline phase as a reference (first gray area). Perturbation was applied after trial 30 and removed after trial 120 (vertical lines). After every 25 trials, a break was provided (dashed vertical lines. **(B)** Final speech adaptation. **(C)** Baseline trial-to-trial variance in F1 frequency. Bar heights and error bars represent robust means and standard deviations, respectively. Dots show individual participant scores. Analysis based on the data from 50 young adults, 79 older adults and 28 older-old adults.

At the end of the adaptation phase (last 30 trials), the level of adaptation, which is a cerebellar-specific measure, was slightly higher in older adults (28.74 ± 38.28 Mels; The mel scale converts physical frequency in hertz into a perceptual pitch scale) than in young adults (26.18 ± 33.33 Mels, Figure 14B). The older-old group exhibited observable lower adaptation (18.32 ± 35.96 Mels). Due to the high inter-individual variability within the groups, these differences did not reach significance (F(2,154) = 1.23, *p* = 0.25). Indeed, F1 frequencies displayed considerable trial-to-trial noise during the adaptation phase, particularly in the older-old adults (young: 25.18 ± 8.11 Mels; older: 25.92 ± 7.70 Mels; older-old: 36.75 ± 11.91 Mels), warranting caution when interpreting potential age-related effects on speech adaptation. Nevertheless, this analysis provides evidence that speech adaptation does not decline with aging (young vs. older: BF = 0.128) but the evidence is not strong for the older-old adults given the intersubject variability (young vs. older-old: BF=0.448).

Baseline F1 trial-to trial variability was taken as a general sensorimotor measure, as voice instability was in older adults has been associated with multiple factors including laryngeal physiology and structural changes in multiple cerebral (motor) regions (Hu et al., 2023). Baseline F1 variability showed a main effect of age on speech control (F(2,154) = 19.4, p < 0.001; Figure 14C). Young adults showed the least variability (19.71 ± 6.91 Mels), comparable to older adults (21.41 ± 7.32 Mels; t(127) = 0.97, p = 0.33, d = 0.18, BF = 0.913). Speech variability in the older-old adults was considerably higher compared to the other age groups (30.57 ± 11.71 Mels; vs. young: t(76) = 4.74, p < 0.001, d = 1.12, BF > 1000; vs. older: t(105)= 4.29, p <0.001, d = 0.94).

Together, these findings indicate that older adults were not impaired in implicit speech adaptation and exhibit only a modest increase in baseline variability, whereas older-old adults show markedly greater speech variability, suggesting emerging sensorimotor control deficits that may contribute to their reduced and more variable adaptation performance.

### Mental rotation

Finally, we went beyond the motor domain by including a mental rotation task, as cerebellar patients have also been shown to exhibit impairments in this ability, which relies on cognitive processes (McDougle et al., 2022). In this task, participants judged whether a rotated letter was presented in a normal or mirrored orientation. Participants from all age groups demonstrated high accuracy in the correctness of their responses (young: 93.24% ± 5.19%, older: 93.48% ± 5.08%, older-old: 81.11% ± 9.60%). Reaction times, calculated from correct responses, increased with larger rotation angles across all age groups (Fig.15A). The slope of the regression line between rotation angles and reaction times indicates the mental rotation pace and serves as a cerebellar-specific measure. The slope varied significantly between age groups (F(2,157) = 7.53, p < 0.001; Fig.15B). Young adults exhibited a reaction time slope (2.62 ± 1.05 ms/°) that is comparable to that of older adults (2.91 ± 1.10 ms/deg; t(128) = 1.19, p = 0.23, d = 0.22, BF = 1.735). In contrast, the older-old group demonstrated a significantly steeper slope (3.6 ± 1.17 ms/°), indicating slower mental rotation compared to both young adults (t(78) = 3.27, p = 0.001, d = 0.75, BF = 746.1) and older adults (t(108) = 2.52, p = 0.01, d = 0.54).

Choice reaction time from baseline trials (0° rotation), which did not involve mental rotation but primarily requires visuospatial attention (Tuch et al., 2005), was used as a general sensorimotor measure. Significant age group differences were observed in baseline reaction times (F(2, 157) = 77.7, p < 0.001; Fig.15C). Young adults demonstrated the shortest reaction time (0.78 ± 0.13 s), while older adults showed significantly longer reaction time (0.97 ± 0.18 s; t(128) = 4.77, p < 0.001, d = 0.86, BF > 1000). Older-old adults (1.4 ± 0.28 s) exhibited the longest reaction time, responding significantly later than both young adults (t(78) = 9.23, p < 0.001, d = 2.84, BF > 1000) and older adults (t(108) = 9.23, p< 0.001, d = 1.98).

These findings suggest that age-related declines in mental rotation pace are accompanied by longer choice reaction times. Although we characterized choice reaction time as a general sensorimotor measure, McDougle and colleagues (2022) reported significant baseline impairments in cerebellar patients. This highlights the importance of considering cerebellar contributions to sensorimotor processes.

**Figure 15.**
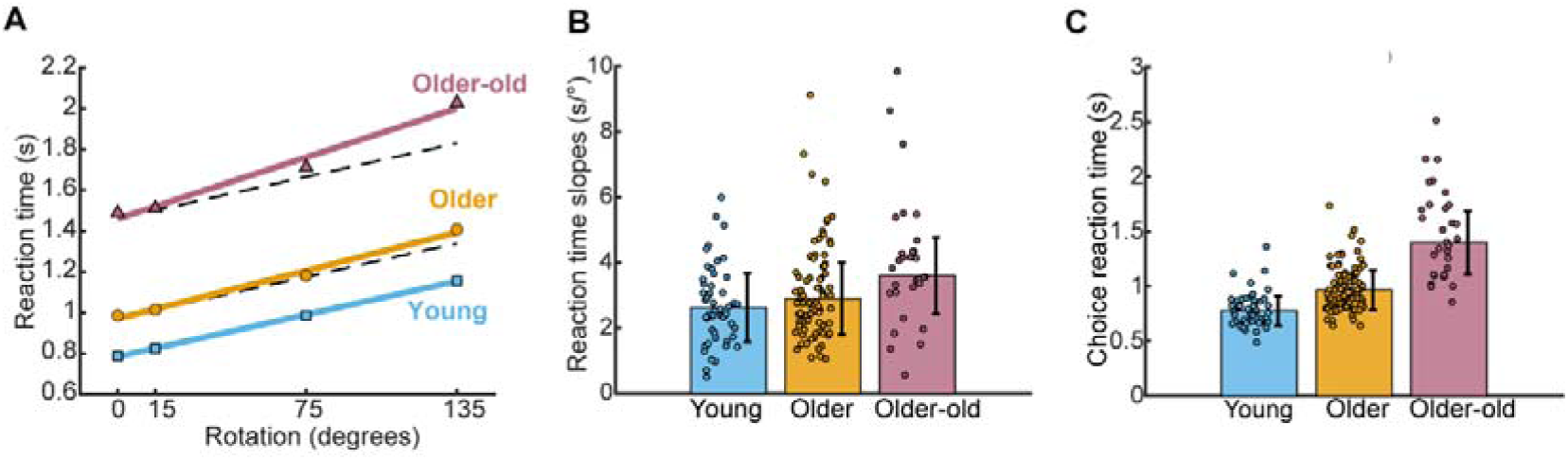
Results from the mental rotation task. **(A)** Reaction times at different rotation angles (symbols) with corresponding regression lines. The black dashed lines represent the regression line of the young adults, for visual comparison with those of older and older-old adults. **(B)** Slopes of the reaction time regression lines, representing mental rotation speed. Steeper slopes indicate slower mental rotation performance. **(C)** Choice reaction time during baseline trials, when no mental rotation was required. Bar heights and error bars represent robust means and standard deviations, respectively. Dots show individual participant scores. Analysis based on the data from 50 young adults, 80 older adults and 30 older-old adults.

### MRI analysis

To quantify age-related changes in brain structure we analyzed MR images and assessed gray and white matter volumes in our sample.

We found no evidence that total intracranial volumes differed across age groups (F(2,143) = 0.10, p = 0.90). The gray matter volume in the cerebellum, expressed as a percentage of the total intracranial volume, differed across age groups (F(2,143) = 16.3, p < 0.001; Fig.16A). Young adults showed the largest cerebellar gray matter volume (6.03% ± 0.36%), with smaller volume in the older adults (5.75% ± 0.41%; t(125) = 3.2, p = 0.002, d = 0.8). In the older-old participants, cerebellar gray matter volume was smallest (5.49% ± 0.34%; vs. young: t(66) = 4.21, p < 0.001, d = 1.47; vs. older: t(96) = 2.99, p = 0.02, d = 0.62).

A similar pattern of differences between age groups was observed in cerebellar white matter volume (F(2,143) = 9.99, p < 0.001; Fig.16B). Young adults had the largest cerebellar white matter volume (2.39% ± 0.15%), which was slightly, but not significantly, larger than that of older adults (2.32% ± 0.22%; t(126) = 1.76, p = 0.08, d = 0.52). In contrast, older-old adults exhibited significantly smaller cerebellar white matter volume (2.17% ± 0.15%) compared to both young (t(66) = 3.45, p < 0.001, d = 1.62) and older adults (t(96) = 2.58, p = 0.01, d = 0.7).

To evaluate whether the average cerebellar volumes of our sample align with previously reported data of a larger reference sample with continuous age data, we analyzed cerebellar gray and white matter volumes on images from the Cam-CAN dataset (n = 653, age range: 19–89 years, mean age: 54.82 ± 18.59 years). Cerebellar gray matter volumes in our sample fell within the range of the Cam-CAN study for participants of comparable age, although cerebellar gray matter volumes in the older and older-old adults in our sample were slightly higher. Moreover, Cam-CAN cerebellar gray matter volumes showed a decline with age (quadratic model: gray matter volume = −1.03 · 10^-4^ · *age*^2^ – 0.0081 · *age +* 6.27), mirroring the age-group differences observed in our sample (Fig.16C).

Cerebellar white matter volumes in our sample tended to fall toward the lower end of the volume range observed in Cam-CAN participants of comparable age. The age-related decline in cerebellar white matter volume with age from the Cam-CAN data (white matter volume = −8.09 · 10^-5^ · *age*^2^ – 0.0028 · *age +* 2.65), mirrors the age-group differences from our sample (Fig.16D). Differences across Cam-CAN cerebellar gray and white matter results and our results may be attributable to differences in MRI acquisition sequences, which could have influenced tissue segmentation. Nevertheless, the observed differences between age groups, when compared to the trends in the Cam-CAN dataset, suggest that cerebellar volumes in our older and older-old adults may be less reduced.

**Figure 16.**
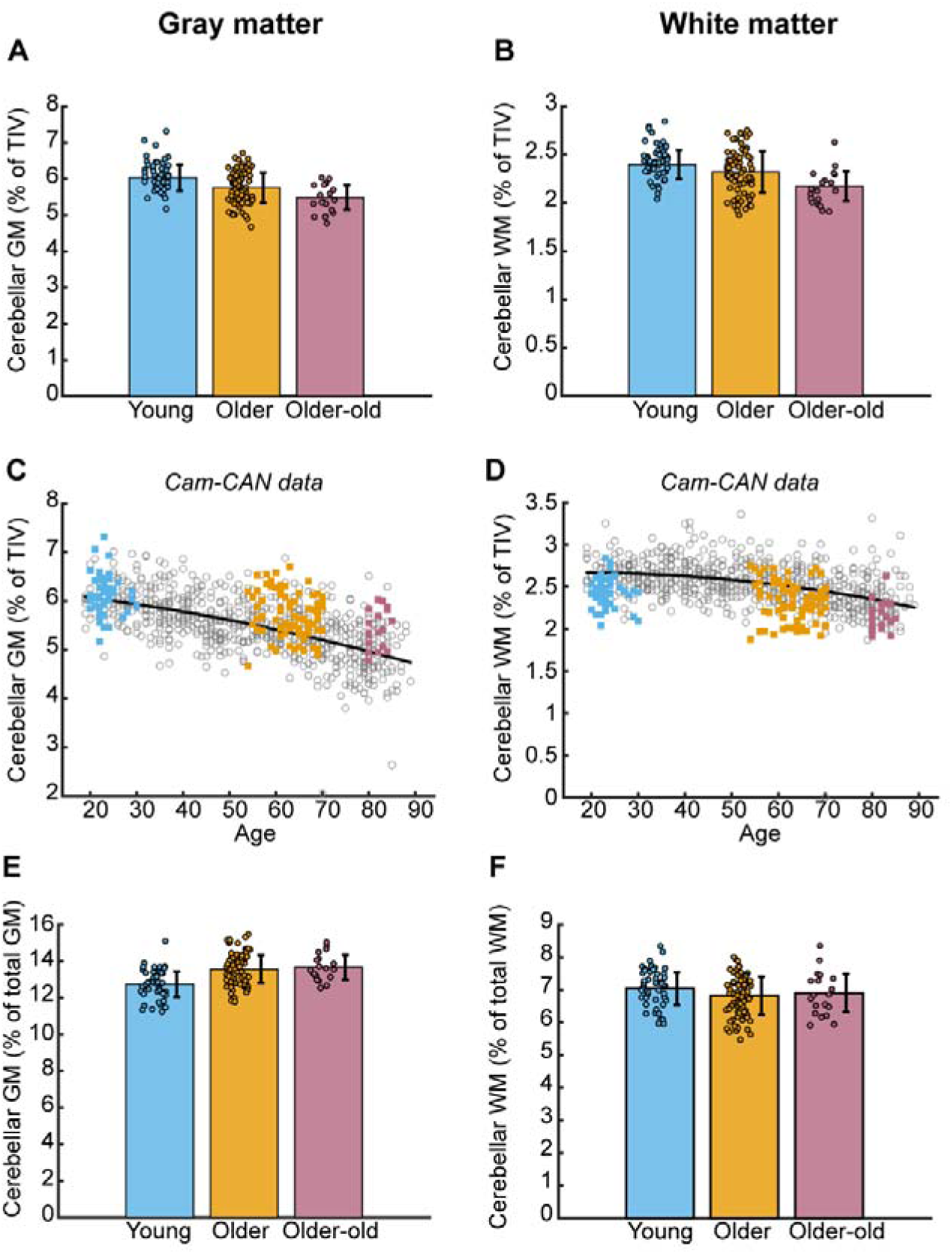
Cerebellar structural measure. The left side of the figure represents our findings in gray matter volumetric results and the right side the white matter results, illustrated with the gray and white matter segments of a young participant**. (A and B)** Cerebellar gray and white matter as percentages of the TIV. **(C and D)** Our results (colored squares) compared to the Cam-CAN dataset. The gray circles indicate the individual data points from the Cam-CAN participants, while the black lines illustrate the quadratic age-related declines in both gray and white matter. **(E and F)** Cerebellar gray and white matter volumes as a percentage of the total gray matter and total white matter, respectively. Bar heights and error bars represent robust means and standard deviations, respectively. Dots show individual participant volumes.

Next, we analyzed the percentages of cerebellar gray and white matter volumes to total brain gray and white matter volumes to determine whether cerebellar gray and white matter reduction was greater, lesser, or comparable to that of the total brain gray and white matter reduction with age. The relative gray matter volume, which is obtained by normalizing the cerebellar gray matter volume by the total brain gray matter volume, increased with age (F(2, 143) = 9.92, p < 0.001; Fig.16E). Here, the young adults exhibited the lowest relative cerebellar volumes (12.74% ± 0.70% of total GM). Relative cerebellar gray matter volumes were markedly higher in the older adults (13.57% ± 0.77% of total GM; t(125) = 4.99, p < 0.001, d = 0.93), being similar to the older-old adults (13.67% ± 0.68% of total GM; vs. young: t(66) = 3.79, p < 0.001, d = 1.04; vs. older: t(96) = 0.46, p = 0.64, d = 0.09). This suggests that the gray matter volume of the cerebellum is decreased less in relation to other parts of the brain in older adults and older-old adults.

We found the opposite result for the relative cerebellar white matter volume, which decreased at older age (F(2,143) = 3.26, p = 0.02; Fig.16F). The relative cerebellar white matter was highest in young adults (7.04% ± 0.58% of total WM). This proportion was slightly, though not significantly, lower in older adults (6.82% ± 0.50% of total WM; *t*(125) = 1.70, *p* = 0.09, *d* = 0.48) and showed a similar pattern in older-old adults (6.91% ± 0.59% of total WM), with no significant differences compared to young (*t*(66) = 0.71, *p* = 0.47, *d* = 0.39) or older adults (*t*(96) = 0.49, *p* = 0.62, *d* = 0.05). These lower relative cerebellar white matter volumes in older groups suggest that cerebellar white matter may decline more in relation to other parts of the brain in older and older-old adults.

All scans had good to excellent image quality (IQR linear rating scale values < 3), however, an effect of age-group on IQR was found (F(2,144) = 20.2, p < 0.001). Therefore IQR was used as a covariate in the MR analysis described above. Images of young adults (mean IQR: 2.04 ± 0.20) had significantly lower quality compared to older adults (1.87 ± 0.13; t(125) = 5.50, p < 0.001, d = 1.00) and to older-old adults (1.81 ± 0.11; t(66) = 4.99, p < 0.001, d = 1.34). No significant difference in image quality was found between older and older-old adults (t(96) = 1.36, p = 0.18, d = 0.35).

### Effect size comparisons and Bayesian Factors across behavioral and structural measures

Based on the previously described analyses, we compiled the effect sizes of all tests comparing older and older-old adults to young adults (Fig.17). We categorized the outcomes into behavioral cerebellar-specific measures, general sensorimotor outcomes, and structural data of the cerebellum.

**Figure 17.**
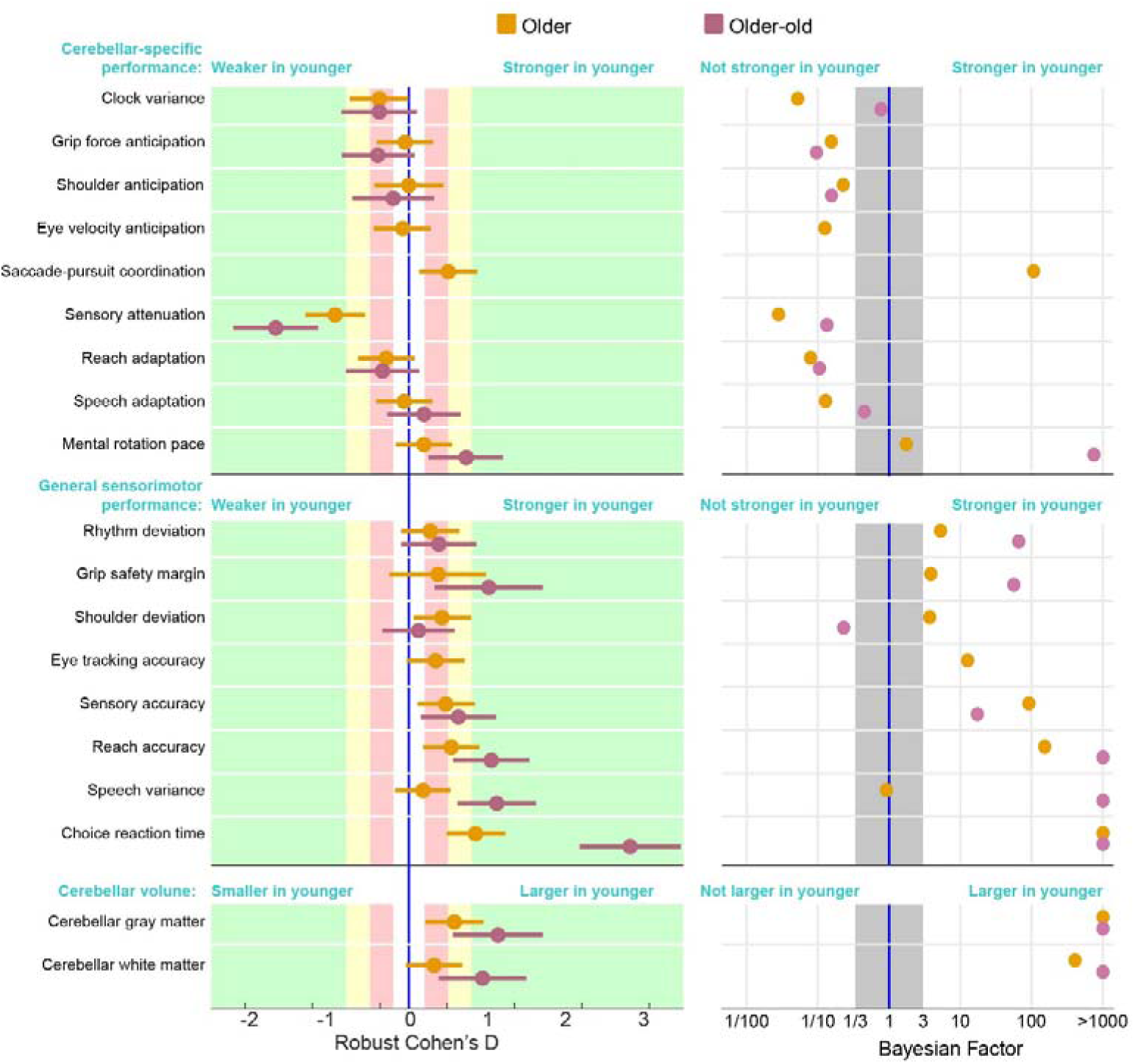
Overview of effect sizes and Bayesian Factors across behavioral and structural measures. Left panel: Each dot represents the effect size from statistical comparisons between age groups: young vs. older adults (orange) and young vs. older-old adults (pink). Horizontal error bars indicate the 95% confidence intervals. Positive values reflect better performance or larger brain volumes in the older groups relative to young adults, while negative values indicate poorer performance or smaller volumes. Cohen’s D effect sizes are interpreted as follows: no effect (<0.2, white), small effect (0.2–0.5, red), medium effect (0.5–0.8, yellow), and large effect (>0.8, green). Right panel: each dot represents the Bayesian factor from statistical comparisons between age groups: young vs. older adults (orange) and young vs. older-old adults (pink). Left from the gray area corresponds to the presence of evidence that the data from the young participants are not stronger (for cerebellar specific and general sensorimotor measures) or larger (for cerebellar volumes) than those of the older participants. On the right side, there is evidence that the performance of the young participants is stronger or that the cerebellar volumes of the young participants are larger than those from the older groups. The gray area represents the area of anecdotal evidence where no conclusion about an age-related effect can be drawn.

Regarding cerebellar-specific measures, older adults showed no decline in motor control tasks and only a slight decline in the cognitive task (mental rotation). Although the effects across most tasks were small, there was a slight overall indication of enhanced cerebellar-specific performance. The Bayesian factor analysis revealed that this data support the fact that cerebellar specific measures are not weaker in older participant groups than in the young group for the majority of the tests that were performed. In contrast, the older adults exhibited slight to moderate declines in general sensorimotor performance. The corresponding Bayes factors support this conclusion. With the exception of shoulder deviation, this decline was even more pronounced in older-old adults. In both age groups, the most substantial decline was observed in choice reaction time, a measure that is arguably more cognitive than purely sensorimotor. The overview reveals a notable decline in cerebellar structure in older adults, with an even greater decline in older-old adults. It shows that the tasks were not too simple as participants from the older groups exhibit a clear decline in general performance. Yet, the cerebellar specific measures did not. These results indicate that while cerebellar structure is declined in older age groups, cerebellar-specific motor control remains largely unaffected by this decline, whereas general sensorimotor and cognitive functions showed varying degrees of age-related deterioration.

Split-half reliability estimates indicated that, although reliability sometimes differed between cerebellar-specific and general sensorimotor measures derived from the same tasks, there was no consistent pattern of greater variability in one measure type relative to the other (Supplementary Table 2). Accordingly, the absence of age-related differences in cerebellar-specific measures is unlikely to be attributable to increased measurement variability.

## Discussion

In this study, we investigated cerebellar motor functions using a series of tasks designed to engage distinct cerebellar processes, ranging from motor timing to mental rotation and speech adaptation. Together, these tasks spanned a broad spectrum of the cerebellum’s known functional domains. Across all these tasks, most outcomes linked to cerebellar function did not show evidence of decline in older age, with only two exceptions: saccade-pursuit coordination and mental rotation. These results challenge the prevailing view that age-related structural changes of the cerebellum are directly responsible for age-related motor control deficits (Bernard & Seidler, 2014; McElroy et al., 2024; Seidler et al., 2010). This view was based on task outcomes that did not selectively represent cerebellar function such as motor learning (Raz et al., 2000; Woodruff-Pak et al., 2010), inter-joint coordination (Seidler et al., 2002), and balance and gait (Droby et al., 2021). When the cerebellum’s contribution to task performance is more precisely isolated, as was done in the present study, the findings suggest a markedly different conclusion: cerebellar functions appear to remain well preserved in older adults.

Two key aspects of our study lend additional strength to this conclusion. First, the absence of age-related differences in cerebellar function was largely confirmed in individuals over 80 years old, an age group older than mostly involved in previous studies reporting preserved cerebellar function, compared to young adults (Arleo et al., 2023). Second, our study is the first to administer the entire task battery to the same group of participants, whereas previous studies typically examined only one of the tasks in isolation (Duchek et al., 1994; Heuer & Hegele, 2008; Parthasharathy et al., 2022; Vandevoorde & Orban De Xivry, 2020; Wolpe et al., 2016). This task battery minimizes task-specific effects, as all tasks share only cerebellar involvement. The consistent absence of age-related differences across tasks thus reflects genuine preservation of most cerebellar functions.

While the selective recruitment of high-performing older adults could be a potential bias, the absence of age-related deterioration in cerebellar function cannot be attributed to this, for two reasons. First, older and older-old participants exhibited clearly reduced cerebellar volumes compared to young adults This structural reduction in older adults was similar to the age-related structural changes reported in other studies studies (Hoogendam et al., 2012; Luft, 1999; Walhovd et al., 2005), and to age effects the Cam-CAN data (Shafto et al., 2014; Taylor et al., 2017), with more pronounced degeneration in participants over 80 years old. Second, we did observe a clear age-related reduction of numerous general sensorimotor measures such as grip force efficiency, sensory accuracy, and movement accuracy (Gilles & Wing, 2003; Wilson et al., 2021; Wolpe et al., 2016). That is, while a cerebellar reserve might exist, it is sufficient to preserve cerebellar function but not all aspects of sensorimotor function.

### Cortical compensation or cerebellar reserve?

The preservation of cerebellar motor functions despite structural degeneration may result from cortical compensation or intrinsic properties of cerebellar processing. It is well established that there is an increase in cortical activation in older adults during motor tasks (Heuninckx et al., 2008; Mattay et al., 2002; T. Wu & Hallett, 2005), which has led to the hypothesis that some cortical areas are recruited to compensate for cerebellar deficits (Marchese et al., 2020). We believe that this is unlikely for two reasons. First, this hypothesis does not align with our findings as such cortical compensation would rather target global sensorimotor task performance instead of cerebellar specific outcomes. Given that sensorimotor performance declined despite the preservation of cerebellar motor functions in our study, it is unlikely that cortical sensorimotor regions play a large role in the cerebellar reserve. Second, the cerebellar circuit is highly specific (Albus, 1971; Ito, 1970), and its function cannot be readily taken over by other brain regions. For instance, patients born without a cerebellum exhibit large motor impairments that cannot be compensated by other brain areas (Leck & Pickett, 2022). Ultimately, however, clarifying the role of cortical compensation in aging will require more comprehensive data, such as task-related functional connectivity between cortical areas and the cerebellum, with careful distinctions between motor and cognitive functions and their corresponding networks.

Several previous studies have already proposed the existence of a cerebellar reserve and the potential neural mechanism behind it (Arleo et al., 2023; Mitoma et al., 2020, 2021) with the idea that the cerebellum is capable of compensating for its structural loss. Cerebellar neuronal circuits can reorganize to counteract local damage due to cerebellar pathological injury (Hillman & Chen, 1985; Mitoma et al., 2021). Similar compensatory mechanisms could also mitigate age-related structural degeneration. A simulation study focusing on the vestibulo-ocular reflex suggested that the preservation of cerebellar-dependent functions after age-related structural changes depends on long-term spike-dependent plasticity of parallel fibers and on the intrinsic plasticity of Purkinje cell synapses (Luque et al., 2022). Further, local cerebellar compensation is compatible with the observation that cerebellar hyperactivity is associated with stable performance in older adults during a predictive motor timing task (Filip et al., 2019).

Given that the cerebellum exhibited a relatively less pronounced structural decline compared to other brain regions as shown here and in another previous study (Taki et al., 2011), it seems more plausible that the cerebellum might compensate for deficits caused by structural changes in other areas rather than vice-versa. Age-related gray and white matter degeneration is usually faster in frontotemporal regions and subcortical regions, including the hippocampus, amygdala and thalamus than in the cerebellum (Fjell et al., 2013; Giorgio et al., 2010; Jernigan et al., 2001; Neufeld et al., 2022). Although this does not directly indicate functional implications, it suggests that cortical regions are less likely to compensate for cerebellar loss when they exhibit more severe degeneration.

Importantly, the relative preservation of cerebellar structure compared to other systems may itself contribute to the maintained cerebellar function observed in older age. Even if structural decline is present, the fact that it progresses more slowly than in many cortical and subcortical regions suggests that a form of structural reserve remains available in the cerebellum. This structural reserve could underlie the continued efficiency of cerebellar circuits and support their capacity to sustain motor functions across aging.

Despite the fact that age-related structural decline in the cerebellum is less pronounced than in the cortex, cerebellar atrophy is clear both in the older and the older-old adults. The preservation of cerebellar motor functions in older age strongly supports the existence of a functional reserve within the cerebellum, which warrants further investigation through the relationship between task performance and corresponding cerebellar regions, which is beyond the scope of the present study, but investigated in more details in another paper (De Witte et al., 2026).

Subtle improvements in cerebellar function suggest it may even compensate for deficits in other sensorimotor functions. For example, the pronounced increase in sensory attenuation (Parthasharathy et al., 2022; Wolpe et al., 2016) could compensate for an age-related decline in sensory accuracy (Wolpe et al., 2016). Similar compensation strategies could explain earlier grip force anticipation observed in our data and the increased motor adaptation (Cisneros et al., 2024; Vandevoorde & Orban De Xivry, 2019). This would further elevate the concept of a cerebellar reserve, suggesting that the cerebellum is not only resilient to its structural decline but also compensates for functional deficits arising from other sources. However, this remains speculative, as the compensations were subtle.

### Variability across cerebellar functions

While we did not observe age-related changes in most cerebellar-specific outcomes, even not in adults above 80 years old, two measures showed age-related changes. First, the coordination between saccades and pursuit systems (Collins & Barnes, 2006; Orban De Xivry et al., 2006) was less precise in older adults than in younger ones. Although there is no data on the effect of cerebellar patients in this task, it is likely that this coordination relies on efferent copies from the saccadic and smooth pursuit systems given the absence of visual signals to drive this coordination (Orban De Xivry et al., 2006). While cerebellar patients cannot adjust their control of saccade amplitude in function of the variability of the peak velocity, older adults can (Xu-Wilson et al., 2009). This suggests that such fine control of saccade amplitude could be cerebellar-dependent. However, other brain regions such as the frontal eye fields (Barborica & Ferrera, 2003) are also involved in this task. Further research is needed to clarify whether the age-related changes in eye movement coordination reflect a deficit in cerebellar function, in broader oculomotor system function, or in another brain region.

Furthermore, our data show that the ability to mentally rotate objects also declines with age. The time it takes to mentally rotate an object increases linearly with the size of the rotation (Shepard & Metzler, 1971) and this continuous mental transformation of the objects occurs more slowly in patients with cerebellar degeneration (McDougle et al., 2022) and in our participants from the older-old group (and, to a lesser extent, from the older group). Either cerebellar cognitive functions are not as well preserved as motor functions or the observed decline in mental rotation ability may be task-specific rather than indicative of a broader cognitive deficit. Future studies employing a more comprehensive test battery that includes a wider range of cerebellar cognitive functions are needed to clarify this issue.

### Definition of cerebellar functions

A major limitation of the current study is the attribution of specific functions to the cerebellum. Our selection of cerebellar functions was based on well-documented deficits in cerebellar patients and, for two outcomes only, on the theoretical framework of cerebellar predictive properties. Cerebellar patients and older adults show overlapping structural loss in the anterior lobes and posterior motor areas, regions associated with motor function. However, degeneration in cerebellar patients is more pronounced in the posterior lobules, which are linked to cognition and higher-order motor functions (Hulst et al., 2015). This pattern suggests that age-related cerebellar degeneration primarily results in motor deficits resembling those observed in patients.

The cerebellum operates within a broader network alongside cortical regions to facilitate motor functions (Shahshahani et al., 2024). Consequently, attributing a single function to the cerebellum is challenging, given its strong connections with non-cerebellar systems. To address this, we distinguished key cerebellar functions from other sensorimotor processes, which are highly important for task performance but do not have the cerebellum as the main driver. It is likely that cerebellar function influences some of our general sensorimotor outcomes such as the safety margin in the grip force task or the reaction time in the mental rotation task, as both of these are increased in cerebellar patients (McDougle et al., 2022; Meindl et al., 2012; Nowak et al., 2005). However, increased mental rotation reaction time are found in Parkinson’s and Huntington’s disease as well, suggesting that the cerebellum contributes to reaction time together with a broader functional network (Jahanshahi et al., 1993). In contrast, outcomes such as the clock variance are impaired in cerebellar patients but not in patients with Parkinson’s disease (Breska & Ivry, 2018).

It is challenging, perhaps even impossible, to fully capture the cerebellar contribution to one task and isolate it from other influencing factors. For this reason, tasks involving deficits commonly observed in cerebellar patients, such as balance and gait problems, were not included. Although these deficits are highly prevalent in older adults, they are also strongly influenced by other factors, including proprioception and muscle strength, which makes it difficult to isolate an outcome that specifically reflects the cerebellum’s contribution (Q. Wang & Fu, 2022). Similarly, no eye-blink conditioning task was included, as it is heavily influenced by cognitive factors such as awareness and arousal, and fear conditioning (LaBar et al., 2004). Previous work has shown that many variables, such as blink reaction time and motor components of the eyeblink reflex, introduce substantial variability in responses at older age (Woodruff-Pak & Jaeger, 1998). In contrast, this study found that only performance on the rhythmic finger-tapping task, similar to what we included in our battery, emerged as a significant predictor of age-related differences in eye-blink conditioning. Furthermore, age-related differences appeared to plateau after early adulthood, with no significant variation in the percentage of correct responses between ages 40 and 80 (Woodruff-Pak & Jaeger, 1998). Practically, the extended duration of the training protocol also makes this task unsuitable for inclusion in a test battery (Winton et al., 2025).

Yet, one of the key strengths of our study in addressing the challenge of assessing cerebellar functions is the use of a comprehensive test battery, incorporating a diverse range of tasks that engage cerebellar functions across different functional networks with a primary focus on motor control. For this reason, we emphasize that our conclusions about cerebellar functions are based on the overall pattern observed across tasks, rather than focusing too narrowly on individual tasks. At the same time, we caution against overgeneralizing cerebellar function across contexts, as it is increasingly evident that the cerebellum can operate in a task-specific manner (Shahshahani et al., 2024).

Similarly, it is important to recognize that general sensorimotor performance is not independent of cerebellar processing. Many broad measures, such as movement accuracy, reaction time, likely reflect contributions from many different brain regions including the cerebellum. As a result, age!zlrelated differences in general sensorimotor performance may emerge from multiple interacting systems rather than cerebellar function alone.

### Conclusion and further research

In conclusion, our study suggests that cerebellar function remains preserved in older adults, even in those above 80 years old, despite significant structural degeneration. While further research is needed to clarify the underlying mechanisms, our findings challenge the notion of direct functional decline due to cerebellar degeneration. This challenges the traditional view of aging as a process of inevitable functional deterioration and opens the door to exploring compensatory mechanisms, such as a cerebellar reserve, that may help maintain functional resilience in older adults.

## Supplementary methods

### I. F1 formant extraction from speech data

Vowels were identified using an amplitude threshold of 0.025 of which the peak with the largest area underneath was selected to contain the vowel. We performed visual and auditory checks on data of all participants to see whether this was correct. For nine participants the original peak selection did not contain the vowel, which could be solved by selecting the first peak instead. For six of them, we additionally increased the amplitude threshold to 0.04 to successfully select the vowel peak.

Trials with short vowel duration (<120 ms) were excluded (1.4% of the trials). Some trials contained multiple sound peaks due to extra articulation of consonants or background noise. In these cases, the largest peak was selected to represent the vowel. If manual checks revealed that this was incorrect for some participants, hand corrections were made. F1 formants were estimated at each time point of the vowel using linear predictive coding.

The mean F1 frequency for each vowel was calculated by averaging the F1 frequencies over the 50 to 100 ms time window following the vowel sound peak onset. This time window was chosen to minimize the influence of the preceding consonant and to exclude any potential within-trial adjustments of the vowel based on auditory feedback. Trials with average F1 frequencies below 250 Mels or above 1000 Mels, indicating incorrect tracking, were excluded from the analysis (1.5% of the trials).

Due to noisy trial-to-trial F1 tracts in some participants, a cluster analysis was conducted to identify outliers in each participant’s F1 data (details in supplementary methods). Two participants with fewer than 10 remaining baseline trials or fewer than 15 remaining adaptation trials after outlier selection were excluded from the final analysis. For the other participants, a total of 713 trials were removed (3.49%).

The F1 tracts were divided into two clusters, which allowed the tracts to be divided into roughly three categories. 1) If the lowest cluster was within three standard deviations of the highest cluster, the F1 tract showed no clear adaptation or outliers. In this case, all F1 values below three standard deviations of the lowest cluster were selected as outliers. 2) If the lowest cluster was more than three standard deviations below highest cluster, while most of the baseline F1 and perturbation F1 values appeared within the highest cluster, the lowest cluster was more likely to represent the outliers and not the adaptation effect. Therefore, all F1 values belonging to the lowest cluster were selected as outliers. 3) If the lowest cluster was more than three standard deviations below highest cluster, with most baseline F1 values in the lowest cluster and most perturbation F1 values in the highest cluster, the lowest cluster was expected to represent baseline values while the highest cluster captured the adapted F1 values. In this case all F1 values below three standard deviations of the lowest cluster (baseline F1), were selected as outliers. This cluster based outlier selection only focused on removing outliers with low F1 values, but not on outliers with high F1 values. First, because this was difficult to quantify due to interference with increased F1 adaptation effects, and second, because we did not notice any high F1 outliers in the data.

## Supplementary results

### I. Inter-joint flexion condition results

During flexion elbow movements, shoulder muscle onset typically occurred before elbow muscle onset (Figure 18A). This timing difference, referred to as shoulder muscle anticipation timing, did not significantly differ between age groups (F(2,95) = 1.94, p = 0.15; Figure 18B). Compared to the shoulder muscle anticipation in young adults (5.41 ± 15.34 ms), the older adults (14.65 ± 19.65 ms) anticipated earlier, while the older-old adults falling between the two groups (9.35 ± 19.27 ms). Although the indications of earlier shoulder muscle anticipation in the older age groups are not statistically meaningful, the results show that shoulder muscle anticipation preceding elbow muscle activation remains intact in older and older-old adults, similar to what we found in the extension condition.

**Figure 18.**
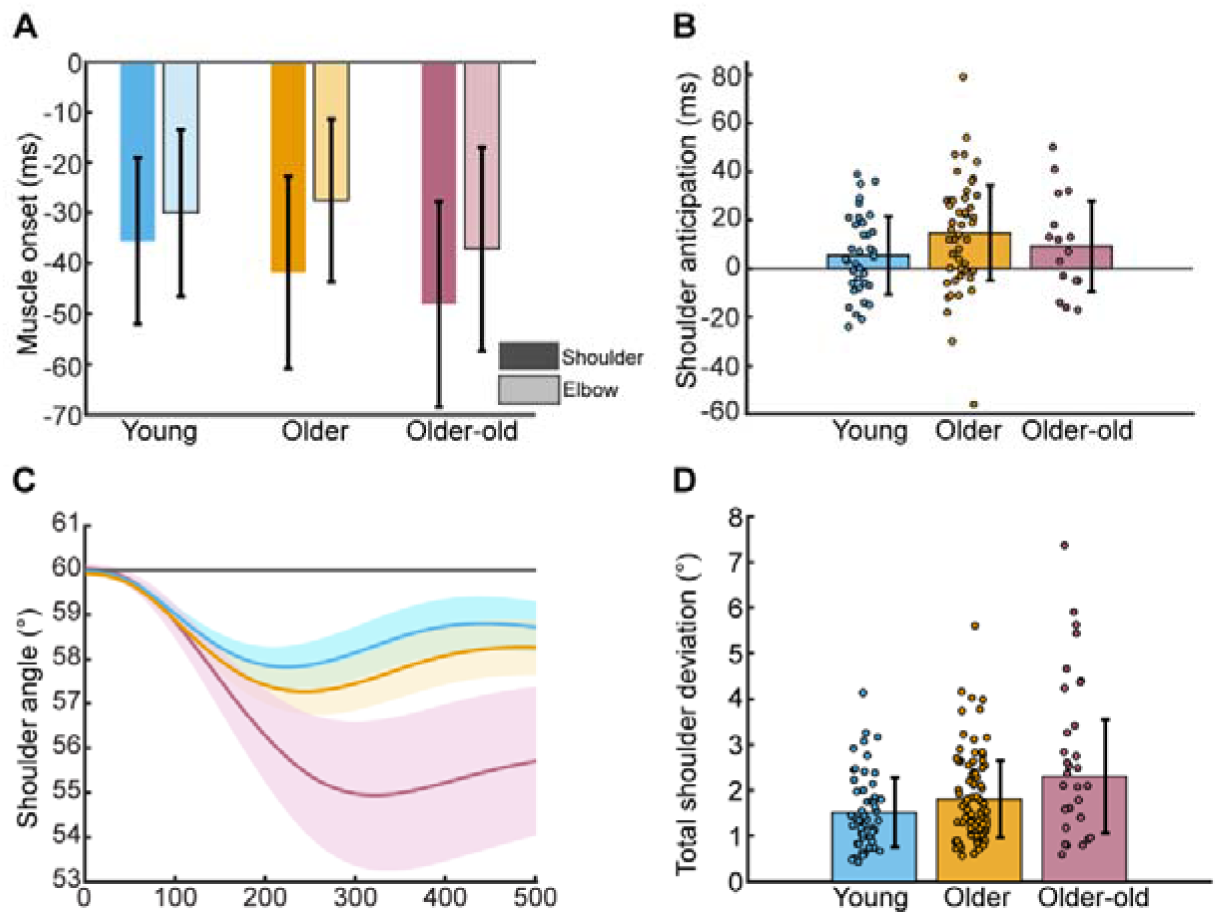
Inter-joint coordination results from flexion condition. **(A)** The timing of the activation of shoulder and elbow muscles before movement onset (at 0 s). Shoulder onset timing is indicated by the darkest color; elbow onset timing is indicated by the lighter color **(B)** Shoulder anticipation timing indicates the time interval between the onset of shoulder muscle activity and the subsequent onset of elbow muscle activity. **(C)** During elbow movement execution, the shoulder deviated from the fixed 60° angle (gray line) across all three age groups. **(D)** Total shoulder deviation during the movement.

Aa a general sensorimotor measure, we assessed the total amount of unintentional shoulder deviation from the fixed angle (60◦; Figure 18C). An effect of age on shoulder deviation was found (F(2,157) = 7.25, p < 0.001; Figure 18D). Young participants showed the least deviation from the fixed shoulder angle (1.52° ± 0.77°), with slightly greater, but insignificant deviation observed in older adults (1.81° ± 0.84°; t(128) = 1.58, p = 0.12, d = 0.29). Total shoulder deviation in older-old adults (2.30° ± 1.24°) was significantly larger than in young adults (t(78) = 3.35, p < 0.001, d = 0.77) and compared to older adults (t(108) = 2.29, p = 0.02, d = 0.49). Although the shoulder deviations pattern in older-old adults looked differently in the extension condition, there is a similarity the pattern of increased unintentional shoulder deviation in the older age groups.

### II. Force matching lever condition results

Given the discrepancy between the absence of overcompensation in young adults from our study and the previous findings (Wolpe et al., 2016), we added the lever condition where the transmission of force between the two fingers was even more direct, increasing the sense of agency (based on Wolpe et al. (2016)). In this condition, participants directly pressed on the lever, using their right finger, to apply force to their left finger. Comparable to force overcompensation in the button condition, normalized lever force overcompensation differed between age groups (F(2, 85) = 4.98, p = 0.009). Young adults exhibited the smallest force overcompensation (0.43 ± 0.34 N, solid line; Figure 19B). Force overcompensation was observably greater in both the older adults (0.82 ± 0.31N, t(57) = 1.59, p = 0.12, d = 0.53) and in older-old adults (0.90 ± 0.29N; t(41) = 2.16, p = 0.03, d = 0.82) compared to young adults, however not significantly. In contrast to the button condition, overcompensation in older-old adults was similar to the older adults (t(107) = 0.92, p = 0.36, d = 0.36). The lack of significance in comparisons involving young adults may be due to the small sample size of young adults (N = 14) in this additional condition.

**Figure 19.**
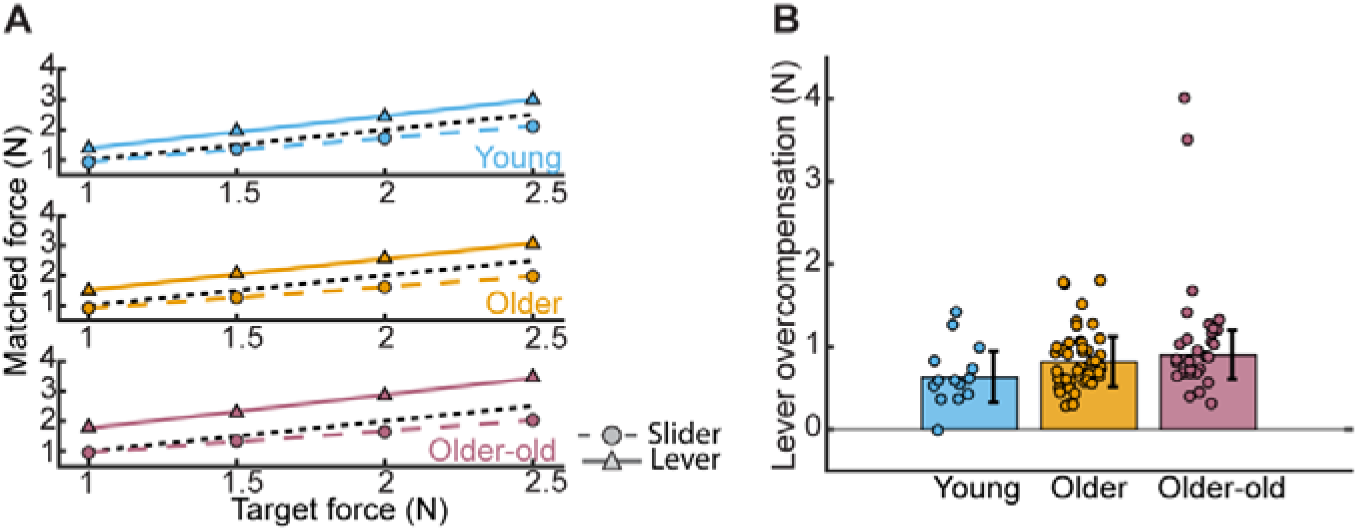
Force matching results from the lever condition. **(A)** Mean linear regression fits of the matched forces against each of the target forces for each age group. The dashed black line is a reference to indicate perfect correspondence between matched and target forces. The dashed colored line with corresponding circles, show the results in the slider condition, which was used as a reference. The solid-colored line and corresponding triangles show the lever condition results. **(B)** Average overcompensation per age group in the lever condition.

### III. Cerebellar gray matter volume with TIV as covariate

We analyzed cerebellar volumes as a percentage of total intracranial volume to account for individual differences in head size. However, upon closer inspection, we observed a strong negative correlation between cerebellar gray matter percentages and total intracranial volume (r = −0.42, p < 0.001), whereas no such relationship was found for white matter. This suggests a potential overcorrection, where individuals with larger total intracranial volumes exhibit disproportionately lower cerebellar gray matter percentages.

Although total intracranial volume did not differ significantly across age groups, we conducted an additional ANCOVA to address this potential bias, incorporating total intracranial volume as a second covariate alongside the IQR values used previously. The inclusion of total intracranial volume as a covariate amplified the observed age-related effects on cerebellar gray matter (F(2,142) = 23.3, p < 0.001). Specifically: Young adults vs. older adults: t(124) = 4.88, p < 0.001, d = 0.80; Young adults vs. older-old adults: t(65) = 5.17, p < 0.001, d = 0.62; Older adults vs. older-old adults: t(95) = 2.64, p = 0.009, d = 0.62. These results indicate that adjusting for total intracranial volume not only addresses potential overcorrection but also strengthens the detected age-related differences in cerebellar gray matter.

### IV. Split-half reliability per task measure

We calculated split-half reliability for each task measure as an index of within-participant response consistency. Reliability estimates were computed separately for each age group to give an overview of measurement stability per measure (Table 2). The grip safety margin measure was excluded from this analysis, as it was derived from mean grip force over a prolonged time interval with few repetitions, making an odd–even trial split inappropriate.

**Table 2.** Split-half reliability per task measure.

| Task | Cerebellar- specific |  |  |  | General sensorimotor |  |  |  |
| --- | --- | --- | --- | --- | --- | --- | --- | --- |
|  | Measure | Young | Older | Older-Old | Measure | Young | Older | Older-Old |
| Rhythmic finger tapping | Clock variance | 0.66 | 0.79 | 0.89 | Rhythm deviation | 0.98 | 0.98 | 0.99 |
| Grip force adjustment | Grip force anticipation | 0.94 | 0.92 | 0.84 | Grip safety margin | - | - | - |
| Inter-joint coordination | Shoulder anticipation | 0.87 | 0.83 | 0.89 | Shoulder deviation | 0.98 | 0.99 | 0.98 |
| Eye movement coordination | Eye velocity anticipation | 0.55 | 0.61 | - | Eye tracking accuracy | 0.97 | 0.97 | - |
|  | Saccade-pursuit coordination | 0.97 | 0.93 | - |  |  |  |  |
| Force matching | Sensory attenuation | 0.94 | 0.94 | 0.96 | Sensory accuracy | 0.73 | 0.84 | 0.86 |
| Reach adaptation | Reach adaptation | 0.96 | 0.98 | 0.98 | Reach accuracy | 0.41 | 0.43 | 0.60 |
| Speech adaptation | Speech adaptation | 0.99 | 0.99 | 0.98 | Speech variance | 0.90 | 0.79 | 0.84 |
| Mental rotation | Mental rotation pace | 0.73 | 0.86 | 0.71 | Choice reaction time | 0.91 | 0.92 | 0.94 |

## Acknowledgement

We thank our master students for helping with data collection.

This work was supported by the Fonds voor Wetenschappelijk Onderzoek Vlaanderen (FWO G095121N).

No conflicts of interest are declared by the authors.

